# Aerial attack strategies of bat-hunting hawks, and the dilution effect of swarming

**DOI:** 10.1101/2020.02.11.942060

**Authors:** Caroline H. Brighton, Lillias Zusi, Kathryn McGowan, Morgan Kinniry, Laura N. Kloepper, Graham K. Taylor

## Abstract

Aggregation behaviors can often reduce predation risk, whether through dilution, confusion, or vigilance effects, but these effects are challenging to measure under natural conditions, involving strong interactions between the behaviors of predators and prey. Here we study aerial predation of massive swarms of Brazilian free-tailed bats *Tadarida brasiliensis* by Swainson’s hawks *Buteo swainsoni*, testing how the behavioral strategies of predator and prey influence catch success and predation risk. The hawks achieved high overall catch success, but they were no more successful against lone bats than against bats flying in column formation. There was therefore no evidence of any net vigilance or confusion effect, and hawks attacking the column benefitted from the opportunity to make several attempted grabs. Even so, the bats’ overall risk of predation was an order of magnitude higher when flying alone. Attacks on lone bats (∼10% of attacks) were greatly overrepresented relative to the proportion of bats classified as flying alone (∼0.2%), so dilution is both necessary and sufficient to explain the higher survival rates of bats flying in the column. From the hawks’ perspective, their odds of catching a bat more than trebled if the attack involved a stoop rather than level flight, or a rolling rather than pitching grab maneuver. These behavioral tactics were independently deployed in nearly three-quarters of all attacks. Hence, whereas the survival rate of a bat depends principally on whether it flies alone or in a group, the catch success of a hawk depends principally on how it maneuvers to attack.

**Lay summary:** Bats emerging by daylight from a massive desert roost are able to minimise their predation risk by maintaining tight column formation, because the hawks that attack them target stragglers disproportionately often. Whereas the predation risk of a bat therefore depends on how it maintains its position within the swarm, the catch success of a hawk depends on how it maneuvers to attack. Catch success is maximised by executing a stooping dive or a rolling grab.

## 1. Background

Flocking, shoaling, and swarming behaviors can all serve to reduce an individual’s predation risk, either by shifting the burden of predation onto others, or by decreasing a predator’s overall hunting efficiency (for reviews see: (Krause 1994; Lehtonen and Jaatinen 2016; Rieucau et al. 2015). The first of these mechanisms is often attributed to a numerical dilution of the risk per attack among the individuals within a group (Foster and Treherne 1981; Morgan and Godin 1985). However, even if the predator only takes one prey item per attack, group members will only enjoy a relative reduction in their predation risk if the predator’s attack rate increases less than proportionally – if at all – with group size (Turner and Pitcher 1986; Wrona and Dixon 1991). The net effect of this is to shift the burden of predation onto lone individuals or smaller groups, in a phenomenon known as attack abatement. Analogous effects apply within a group, with individuals at the periphery expected to suffer higher attack rates than those in the centre (Hamilton 1971). This phenomenon is known as marginal predation, and has been widely observed across taxa (Duffield and Ioannou 2017; Ioannou et al. 2017; Rayor and Uetz 1990) but see (Parrish 1989). The second class of mechanism, involving an outright reduction in the predator’s hunting efficiency, is usually attributed either to the benefits of shared vigilance (Lima 1995), or to the confusion occurring when the presence of multiple prey makes it harder for a predator to target any one individual (Duffield and Ioannou 2017; Landeau and Terborgh 1986; Quinn and Cresswell 2006). Confusion may well have an effect on the outcome of a directed chase, but a predator lunging or plunging into a dense prey aggregation need not be targeting any one individual. In such cases, hunting may be more efficient against a denser group. This is true, for example, of those large pelagic predators such as baleen whales that exploit the shoaling of their prey during engulfment (Cade et al. 2020), and might also be true of raptorial predators that will have more opportunities to capture individual prey items when striking at a dense school or swam (Nottestad and Axelsen 1999).

Understanding the behavior of predators is therefore key to understanding the ecology and evolution of their interactions with prey (Hein et al. 2020; Lima 2002). For example, the group-size dependence of attack frequency and catch success has been found to vary with predator hunting-mode and species in raptors attacking flocks of waders, leading to conflicting selection pressures on group size (Cresswell and Quinn 2010). On a finer scale, the dynamics of collective motion in schooling fish is not only influenced by their direct response to predator behavior (Cade et al. 2020; Handegard et al. 2012; Magurran and Pitcher 1987), but is also governed by attraction and orientation rules that may have evolved to promote the formation of coherent mobile groups that cause cognitive or sensory confusion in predators (Ioannou et al. 2012). Nevertheless, idealized approaches have long prevailed when modelling predator behavior – from mass action kinetics in models of prey encounter (Hutchinson and Waser 2007), to fixed attack rates in models of prey capture (Morrell and Romey 2008). This partly reflects the level of complexity that it has been thought necessary and sufficient to capture in generating ecologically relevant models of predator-prey interactions (Hein et al. 2020). For instance, the potential significance of fine-scale search behavior to the long-run population dynamics of predators and prey has only recently been made clear (Hein and Martin 2020). Likewise, the importance of incorporating physical and physiological constraints into models of fine-scale chase dynamics has also only lately been revealed (Mills et al. 2018; Mills et al. 2019). Above all, the over-idealization of behavior in empirical modelling reflects the extreme difficulty of making repeatable observations of predator-prey interactions in their natural ecological context, rather than in laboratory settings.

Nowhere is the challenge of making field observations clearer than in the case of predators attacking massive three-dimensional prey aggregations. The best examples to date have come from sonar studies of pelagic predators attacking schooling fish (Axelsen et al. 2001; Gerlotto et al. 2006; Handegard et al. 2012; Nottestad and Axelsen 1999; Similä 1997), and videographic studies of aerial predators attacking murmurating birds (Carere et al. 2009; Procaccini et al. 2011; Storms et al. 2019). These sonar imaging studies have successfully related predator attack behavior to shoal size, shape, and density, but have not yet allowed the individual outcomes of these behaviors to be observed (Handegard et al. 2012); videographic studies have been able to record individual outcomes, but have focussed on the dynamics of the prey’s collective motion, rather than the dynamics of the predator’s attack (Carere et al. 2009; Procaccini et al. 2011; Storms et al. 2019). Here we use high-definition videography to analyse the behavioral ecology of predator-prey interactions between Swainson’s hawks (*Buteo swainsoni*) and massive swarms of Brazilian free-tailed bats (*Tadarida brasiliensis*) emerging in column formation from their roost. First, we test whether maintaining column formation confers an adaptive benefit on individual bats by reducing their overall predation risk, and if so whether this benefit is attributable to dilution, confusion, or vigilance effects. Second, we test whether bats are more likely to be captured in attacks from above and behind, as would be expected if vigilance were an important feature of this system, given their forward-facing eyes and sonar (Lima and O’Keefe 2013). Third, given the observed variation in attack maneuvers, we test whether high-speed attacks involving stooping dives and rolling grab maneuvers result in higher catch success than attacks involving level flight and pitching grab maneuvers, as a recent physics-based simulations of raptor attack behaviors predicts (Mills et al. 2018). In testing these hypotheses, we provide the ecological context for future work that will analyse the three-dimensional attack and escape trajectories of observed predator-prey dyads from a biomechanical and algorithmic perspective (see (Brighton and Taylor 2019; Brighton et al. 2017)).

## 2. Methods

We observed Swainson’s hawks hunting Brazilian free-tailed bats at the Jornada Caves, New Mexico, USA. This remote site is located on a lava field, where a collapsed lava tube forms a connected pair of caves (Fig. 1A). The caves are home to a maternal colony of 700,000 to 900,000 individuals (Kloepper et al. 2016), which migrate to the area during their natal season from May to September. The bats roost during the day, emerge before dusk to fly to their feeding grounds, and return individually or in small groups towards dawn. The emerging bats form a dense horizontal column (Fig. 1B) which climbs away from the cave, giving the appearance first of a rising plume of smoke, then of distant clouds as the swarm splits into smaller groups. The local population of Swainson’s hawks (Fig. 1C) hunts the emerging bats daily. The bats are subject to regular predation by great horned owls (*Bubo virginianus*) within the cave mouth, but the only other aerial predation events that we witnessed over three field seasons involved a single peregrine falcon (*Falco peregrinus*) hunting on three consecutive evenings in 2018.

**Figure 1.**
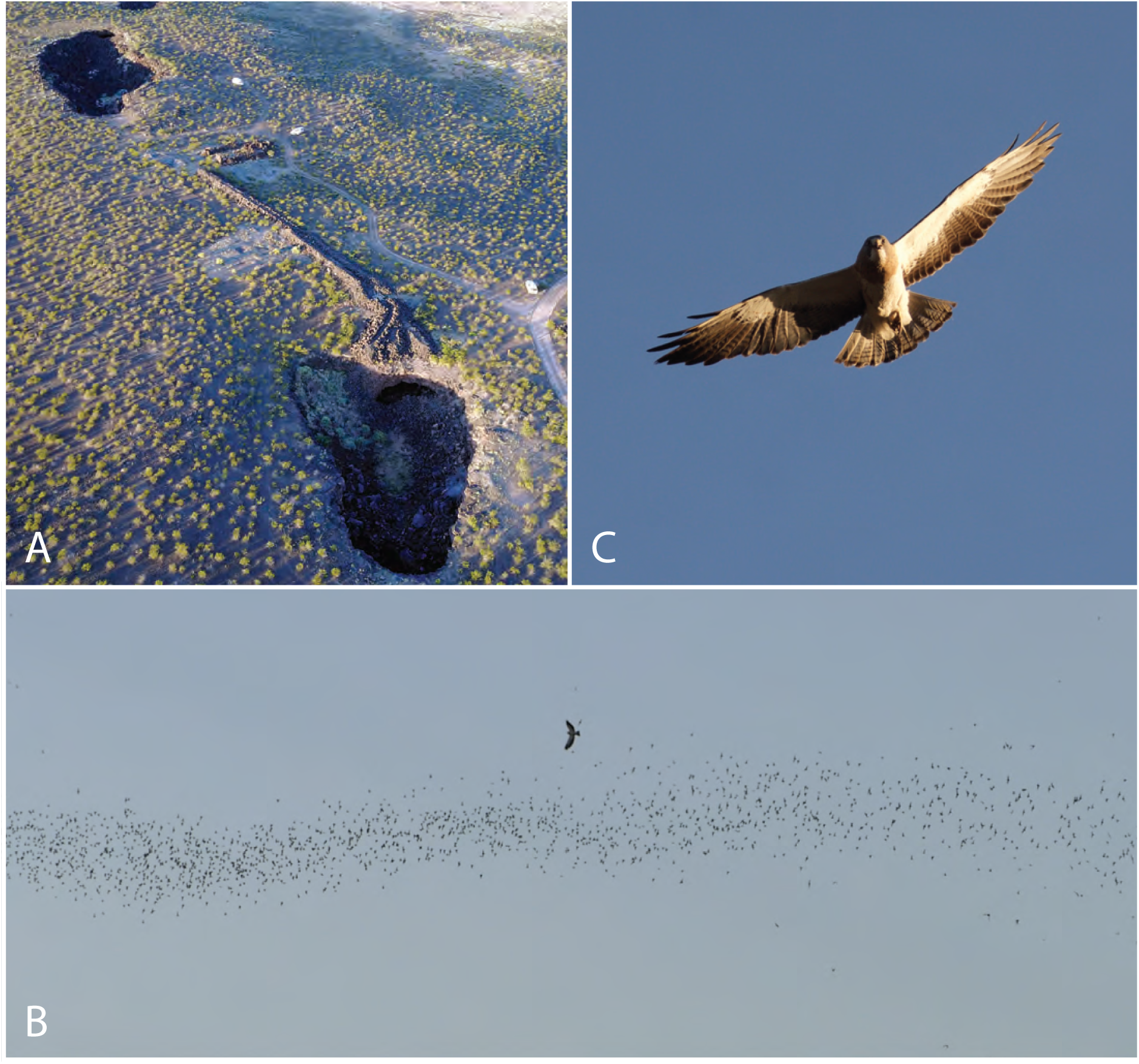
Study overview. (A) Aerial view of the Jornada caves. (B-C) Swainson’s hawk hunting Mexican free-tailed bats at the caves. Note the column formation of the swarm in (B), with only a small proportion of bats flying alone.

### A. Behavioral observations

We systematically recorded the hawks’ hunting behavior on 15 evenings from 01/06/18 to 24/06/18, having routinely witnessed the same behaviors in 2016 and 2017. Emergence began between 18:19h and 19:52h, and hence well before sunset, which was between 20:13h and 20:21h. The timing of the bats’ emergence was quite variable, but the hawks usually appeared within a few minutes of its onset, suggesting that they must have been watching the caves from a distance. The number of hawks present varied throughout the observation period, peaking at ∼20. Each emergence lasted from 10 to 25 mins, depending on the number of bats emerging, and we occasionally observed a second emergence from the same cave. We conducted focal follows using a voice recorder to document real-time observations made through 8×4 binoculars, or using a Lumix DMC-FZ1000/2500 camera (Panasonic Inc., Osaka, Japan; 1920×1080 pixels; 50 fps) recording video for later analysis (see Movie S1). Each observer aimed to document the entire hunting bout of one focal hawk, from its first appearance to its final departure. We began by observing the hawks from makeshift hides, but phased these out as the birds became habituated to our presence. We measured wind speed using a Kestrel 4500 Pocket Weather Tracker (Nielsen-Kellerman, PA, USA), and used the NOAA Solar Calculator (National Oceanic and Atmospheric Administration U.S. Department of Commerce 2018) to determine the time of emergence relative to sunset. We statistically controlled for these environmental variables when testing the behavioral factors affecting capture outcome, on the basis that catch success might be expected to vary with wind speed and light level.

### B.Behavioral classification

The hawks usually made multiple attack passes in a single hunting bout. We categorised each attack pass according to: 1) approach type: level flight, or stooping dive; 2) approach direction: downstream, cross-stream, or upstream relative to bat(s); 3) grab direction: above, beside, or below targeted bat; 4) target type: lone bat, or column; and 5) capture outcome: success or failure; see Table 1, Figs. 2, S1 and Movie S1. A single attack pass could sometimes involve more than one attempted grab if the earlier grab(s) had been unsuccessful, in which case we recorded the direction and outcome of the final grab only. Our estimates of catch success therefore provide an unbiased estimate of the success rate per pass, but not of the success rate per grab – except in the case of attacks on lone bats, which usually involved only one grab. In total, we observed the outcomes of 239 attacks from 64 hunting bouts lasting 2h50m (Fig. 3B), and were able to categorise 202 of these attacks fully (Fig. 3A; Table S2; Data S1).

**Table 1.** System used to classify hawk attack behaviors.

|  |  |
| --- | --- |
| approach type | <u>Level flight</u>: flapping or gliding along a shallow flight path (Fig. 2A)<br><u>Stooping dive</u>: fast descent on tucked wings, along a steep dive path (Fig. 2B) |
| approach direction | <u>Downstream</u>: hawk flying in same direction as bat(s) (Fig. 2C)<br><u>Upstream</u>: hawk flying in opposite direction to bat(s) (Fig. 2C)<br><u>Cross-stream</u>: hawk flying in any other direction relative to bat(s) (Fig. 2C) |
| grab direction | <u>Above</u>: grab initiated from above bat, in a pitch-down maneuver with the legs extended downwards (Fig. 2D)<br><u>Beside</u>: grab initiated from beside bat, in a roll maneuver with the legs extended horizontally (Fig. 2E)<br><u>Below</u>: grab initiated from beneath bat, in a pitch-up maneuver with the legs extended upwards (Fig. 2F) |
| targeting strategy | <u>Lone bat</u>: attack on bat flying >5 body lengths from the edge of the column, or in a different direction to the column (Fig. S2 C-F)<br><u>Column of bats</u>: attack on one or more bats flying within a cohesive group of individuals, all flying in the same general direction (Fig. S2 A,B) |
| capture outcome | <u>Success</u>: bat caught, independent of whether subsequently dropped or eaten<br><u>Failure</u>: bat not caught |

**Figure 2.**
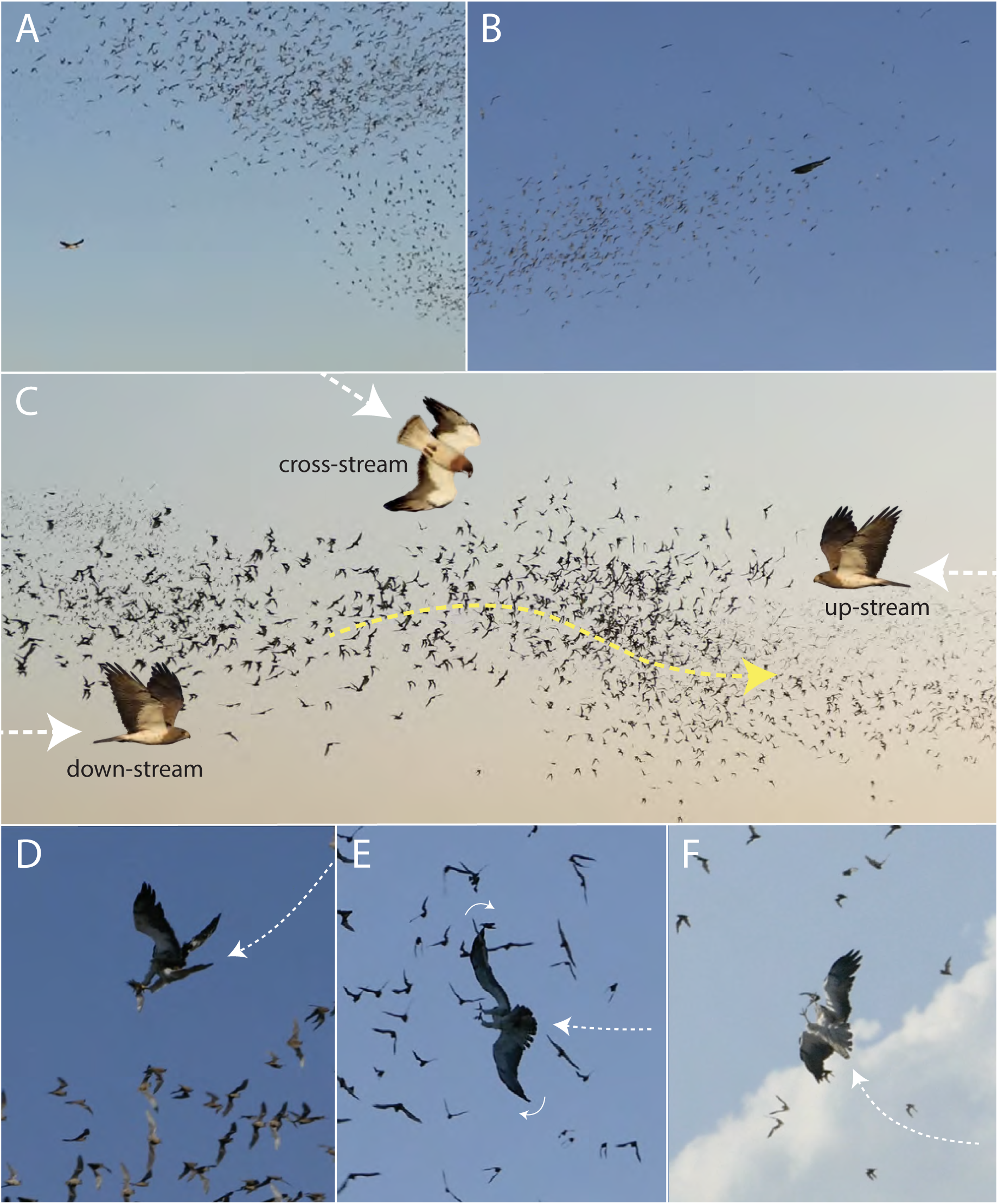
Examples of categorised attack behaviors. (A,B) Approach type, showing: (A) level flapping flight toward the column; (B) stooping dive into column, with tucked wings. (C) Approach direction, with composite image comprising video frame of swarm moving from left to right, superimposed with separate images of Swainson’s hawks to illustrate upstream, downstream, and cross-stream approach. (D-F) Grab direction, with video frames showing: (D) grab from above bat: bird extending feet downwards in a pitch-down maneuver; (E) grab from beside bat: bird extending feet horizontally in a roll maneuver; (F) grab from below bat: bird extending feet upwards in a pitch-up maneuver. White arrows indicate approximate attack trajectory. See Movie S1 for video examples.

**Figure 3.**
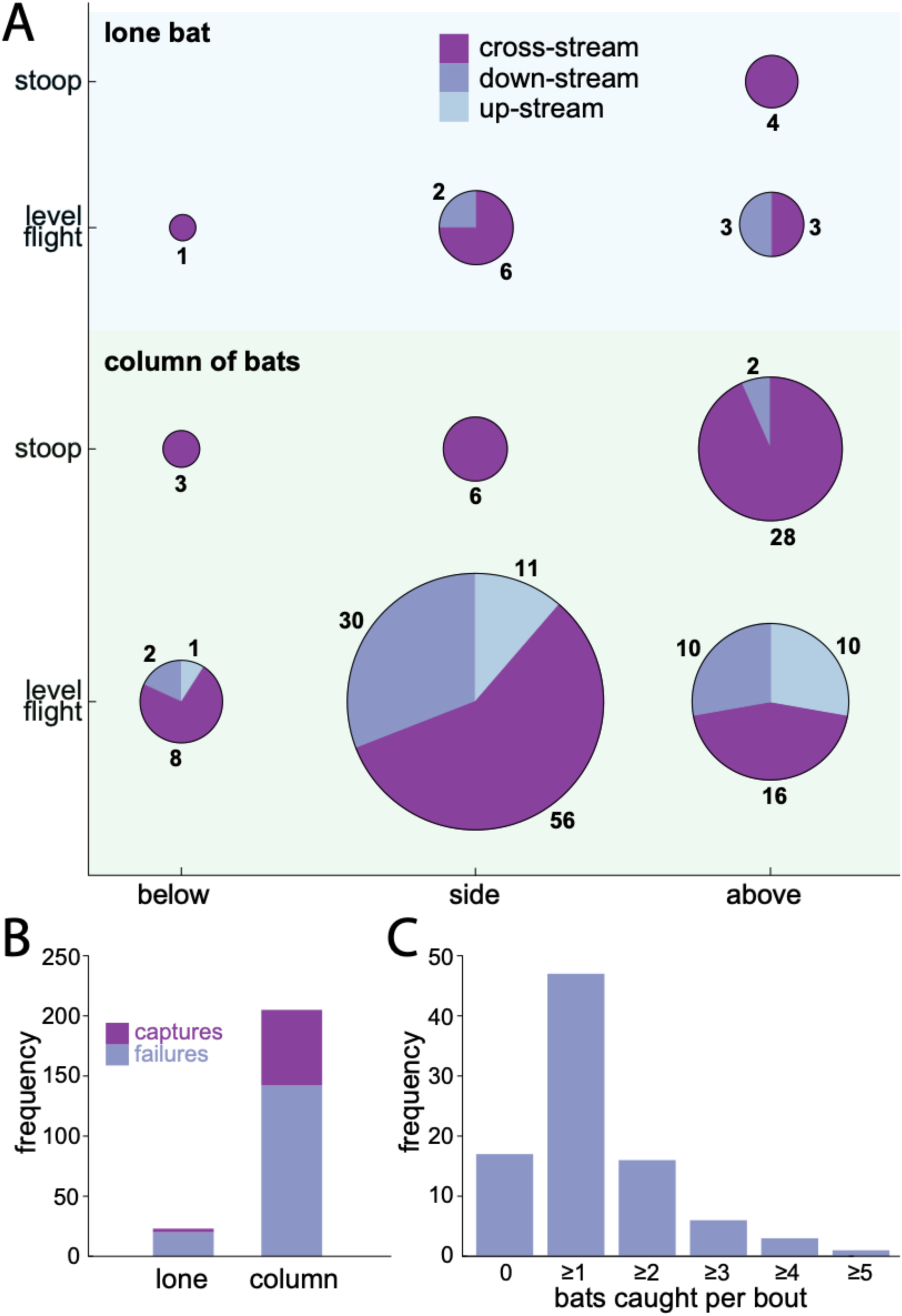
Behavioral strategies of bat-hunting Swainson’s hawks. (A) Pie charts showing frequency of each combination of behaviors for all 202 fully classified attacks; total area of each pie chart proportional to total number of observations shown by each segment. (B) Bar chart showing number of successes and failures for all *n* = 239 attacks, grouped by target type. (C) Frequency distribution of number of bats caught per hunting bout; inset shows typical feeding posture.

Our assessment of target type follows the recommended practice of considering both an absolute distance criterion and a behavioral difference criterion when classifying prey as solitary or grouped (Stankowich 2003). We classified a targeted bat as a lone individual if it appeared to be >5 body lengths away from its nearest neighbor in the two-dimensional field of view of the imaging system. The projective geometry of the imaging system guarantees that any bat for which this criterion holds must be >5 body lengths from its nearest neighbor in three dimensions, so identifying lone bats using this method should not produce any false positives. On the other hand, this method is expected to produce false negatives for bats silhouetted against the column, so we also classified a targeted bat as a lone individual if it was clearly flying at a velocity different to the column. In practice, this behavioral difference criterion proved most useful in identifying stragglers that were flying in close but uncoordinated proximity to another. The application of these criteria is expected to minimise false negatives, but not to eliminate them completely, so we expect that a small fraction of the bats classified as flying in the column will in fact have been flying >5 body lengths from it (see Discussion).

We estimated the swarm dilution factor *D*, defined as *D* = *N*_C_/*N*_L_ where *N*_L_ is the number of bats flying in the column and where *N*_L_ is the number of lone bats, from a sample of 18 frames selected from across the videos (Fig. S2). These frames were chosen as meeting the following requirements: (i) each frame recorded in a separate attack; (ii) camera zoomed out and in focus; (iii) bats near enough to see their wings; and (iv) background composed entirely of sky. Having counted the number of bats meeting the criteria for classification as lone bats in each frame, we then estimated the total number of bats automatically using the count function in Adobe Photoshop CC2019, having binarized each image using a threshold just sufficient to make the background entirely white.

### C. Statistical analysis

We conducted the statistical analysis in *R* version 3.6.1 (RCoreTeam 2019), using the *PropCIs, plyr*, and *boot* packages. The raw data and statistical code are provided in Data S1 and Code S1. As far as possible, the analysis was designed to account for the non-independence of attacks within the same hunting bout, noting that there was no way of identifying individuals to control for the possibility that we had sampled the same individual repeatedly across days. We denote sample proportions for the observed attacks using the notation 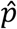, and use standard probability notation to identify the proportion to which the quantity refers. Hence, 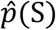 estimates the probability *P*(S) that an attack is successful, 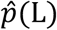 estimates the probability *P*(L) that an attack is on a lone bat, and 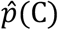(C) estimates the probability *P*(C) that an attack is on a bat flying in the column. Hence, 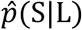 is the proportion of the observed attacks on lone bats that were successful, which estimates the probability *P*(S|L) that an attack on a lone bat proves successful.

As the raw data are counts, we computed a 95% profile-likelihood confidence interval (CI) for the mean attack rate *λ* over all *n* = 64 bouts using an intercept-only quasi-Poisson regression with log bout duration as an offset variable. We computed a 95% profile-likelihood CI for the mean number of bats caught per bout using an intercept-only quasi-Poisson regression with no offset variable. Adding the log number of attacks by bout as an offset variable allowed us to compute a 95% profile-likelihood CI for 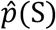, defined as the mean per-attack success rate over all bouts. Hawks with a lower success rate may have made a greater number of attacks, and thus contributed disproportionately to the sample used to calculate 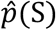. To eliminate this potential bias, we also report 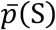, defined as the mean per-attack success rate averaged by hunting bout, together with a bias-corrected and accelerated bootstrap 95% CI computed using stratified resampling over 10^6^ resamples. The CIs that we report for 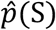 and 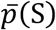 are thereby constructed so as to avoid pseudo-replication within a bout. Other percentages or proportions relating to pooled subsets of the data (e.g. 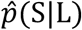, defined as the mean per-attack success rate for all attacks on bats classified as lone bats) were calculated directly, and are reported together with a 95% Wilson score CI. These score CIs may have less than nominal coverage probability because of the non-independence of attacks from the same hunting bout, but we report them in preference to stating only a point estimate. Odds were calculated directly from the relevant proportions and their associated CIs.

We use autoregressive logistic regression for statistical inference relating to the factors influencing capture outcome. The first-order autocorrelation coefficient for the outcomes of the subset of *N* = 175 attacks that were preceded by another from the same hunting bout (*r*_1_ = 0.239) fell well outside the 95% CI for white noise (−0.148, 0.148), indicating the presence of statistically significant runs of successful or unsuccessful attacks. To avoid having to discard the first attack of each bout in our autoregressive logistic regressions, we concatenated the time series data for consecutive hunting bouts. This approach effectively substitutes white noise for the autoregressive term corresponding to the first attack of each bout. Provisional model-order selection using the Akaike Information Criterion (AIC) in a pure autoregressive model supported our use of AR(1) logistic regression to model the factors affecting capture outcome. We also used an AR(1) logistic regression model to test whether other aspects of attack behavior predict target type. We used likelihood ratio tests to assess the significance of the model factors, Wald tests to assess the significance of differences between their levels, and profile-likelihood 95% CIs to quantify the uncertainty in the parameter estimates. Odds ratios from the logistic regressions were computed by exponentiating the logistic regression coefficients.

## 3. Results

### A. Hawks hunting swarming bats achieve high catch success at high intensity

Most of the *n* = 64 hunting bouts that we observed involved multiple attacks (median: 3; 1^st^, 3^rd^ quartiles: 2, 5; maximum: 15). These were made at high intensity, with a mean attack rate of *λ* = 0.0234 s^−1^ over all bouts (CI: 0.0175, 0.0304 s^-1^; dispersion parameter in quasi-Poisson regression: *ϕ* = 4.66; df = 63). Three-quarters of the observed bouts resulted in the capture of at least one bat (75.0%; CI: 63.2%, 84.0%). The hawks invariably consumed their prey on the wing, so were able to catch more than one bat per hunting bout, with a mean catch of 1.16 bats per bout (CI: 0.92, 1.43; dispersion parameter in quasi-Poisson regression: *ϕ* = 0.97; df = 63; Fig. 3C). Bats were caught at a mean per-attack success rate over all bouts of 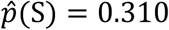 (CI: 0.241, 0.390; dispersion parameter in quasi-Poisson regression: *ϕ* = 1.11; df = 63). This estimate of the hawks’ per-attack success rate equates to the overall proportion of attacks that were successful, so will systematically underestimate the mean per-attack success rate by individual if hawks with a lower success rate make more attacks and thus contribute disproportionately to the sample. Indeed, the mean per-attack success rate averaged by hunting bout was significantly higher at 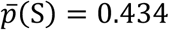 (bootstrap CI: 0.395, 0.475; *n* = 64 bouts), which implies that individuals with a higher success rate made fewer attacks – presumably because they reached satiation sooner.

### B. Flying in the column reduces predation risk in swarming bats

Only 5% of the bats that we observed being caught were classified as lone bats (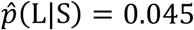 CI: 0.016, 0.125; *n* = 66 captures classified by target type; Fig. 3B). This represents a small proportion of the total catch, but is still many times higher than the proportion of the population meeting the criteria for classification as lone bats (0.2% of >34,000 bats visible in 18 frames; Fig. S2, Table S3). It follows that the predation risk of bats flying alone must have been correspondingly higher than the predation risk of bats flying in the column. In fact, the relative risk of capture for a lone bat is just:

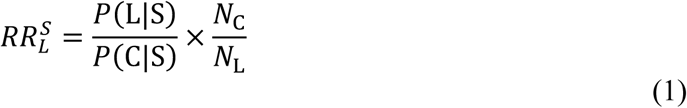

where the first term on the righthand side is the odds *O*_L|S_ that a captured bat flies alone rather than in the swarm (sample odds: *ô*_L|S_ = 0.048; CI: 0.016, 0.143; *n* = 66 captures classified by target type), and where the second term is the ratio of the number of bats flying in the column to the number of bats flying alone, which we will call the swarm dilution factor *D* = *N*_C_/*N*_L_. Hence, since 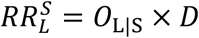, lone bats are expected to experience a higher risk than bats flying in the column 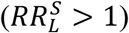 if and only if *D* ≥ 21 (CI: 7, 63). Hence, as we estimate that *D* > 500 (Table S3), it is clear that flying in the column significantly reduces a bat’s predation risk.

### C. Dilution, not vigilance or confusion, reduces predation risk in the column

To identify the mechanism underlying this reduction in predation risk, we may use Bayes’ theorem to rewrite Eq. 1 as:

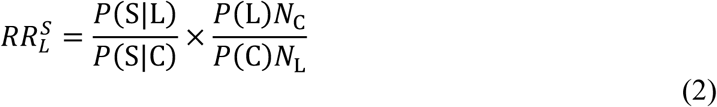

where the first term on the righthand side is the relative catch success of a hawk attacking a lone bat versus a hawk attacking the column. Clearly, this ratio would have to be significantly greater than one for us to conclude that either vigilance or confusion was important in explaining the lower predation risk to which swarming bats were exposed. In fact, there was no significant difference in the hawks’ overall catch success for attacks on lone bats versus attacks on the column (likelihood ratio test in AR(1) logistic regression: *χ*^2^ (1) = 2.582, *p* = 0.108; *n* = 228 attacks categorised by target type), so there is no evidence of any significant vigilance or confusion effect (Fig. 3B). The second term on the righthand side of Eq. 2 represents the combined effects of dilution and avoidance in attack abatement, where *D* = *N*_C_/*N*_L_ is the swarm dilution factor, and where *O*_L_ = *P*(L)/*P*(C) is the odds that a hawk attacks a lone bat (sample odds: *ô*_L_ = 0.112; CI: 0.073, 0.172; *n* = 228 attacks categorised by target type). These combine to determine the relative risk of attack 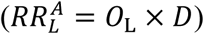, where we require that 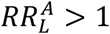 to conclude that attack abatement is occurring, which holds if and only if *D* ≥ 9 (CI: 6, 14). Given that we estimate that *D* > 500 (Table S3), we conclude that dilution is both necessary and sufficient to explain the lower predation risk for bats flying in the column. In other words, lone bats were disproportionately more likely to come under attack than bats flying in the column, albeit that the great majority of attacks were still made against the column.

### D. Hawks attacking the column have multiple opportunities to grab a bat

Although we never observed more than one bat being captured in a single attack, presumably because of the processing that was required, attacks on the column could sometimes involve up to three attempted grabs if the preceding grab(s) had been unsuccessful. Other things being equal, we might therefore have expected to observe a higher success rate per attack against the column. In principle, the expected catch success of an attack involving up to *k* independent grabs is *P*(S)|_*k*_ = 1 − (1 − *q*) ^*k*^, where *q* represents the probability that a given grab proves successful. Treating the observed catch success against lone bats (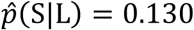; CI: 0.045, 0.321; *n* = 23 attacks on lone bats) as an estimate of *q*, the expected catch success of an attack involving up to *k* = 3 independent grabs, *P*(S)|_*k*=3_ = 0.342, is statistically indistinguishable from the overall catch success for attacks on the column (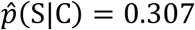; CI: 0.248, 0.374; *n* = 205 attacks on the column). Hence, although we are not in a position to conclude that the hawks were any more or less successful in their attacks against the column than against lone bats (see above), our results are consistent with the possibility that the hawks might have improved their effective success rate when attacking the column by taking multiple opportunities to grab a bat.

### E. Stoops and rolling grab maneuvers are associated with higher catch success

We used AR(1) logistic regression to model how the odds of capture were related to approach type, approach direction, grab direction, target type, wind speed, or time before sunset. Only approach type and grab direction were significant in the full AR(1) model (likelihood ratio tests at *α* = 0.05). A reduced AR(1) model retaining only these two factors was better supported than the full AR(1) model (ΔAIC = 7.2), and was better supported than a pure AR(1) model without any factors (ΔAIC = 6.0), so we are confident of their effect. There was no evidence of any statistically significant interaction between approach type and grab direction (Fig. 4D), as the reduced AR(1) model containing only their main effects was better supported than a model also containing their interaction (ΔAIC = 3.8). The expected odds of capture in the reduced AR(1) model were 3.46 times higher (CI: 1.43, 8.82) in a stoop than in level flight (Wald test: *z* = 2.70, *p* = 0.007), and were 3.45 times higher (CI: 1.54, 8.48) if the bat was grabbed in a rolling maneuver from the side rather than in a pitching maneuver from above (Wald test: *z* = 2.87, *p* = 0.004).

**Figure 4.**
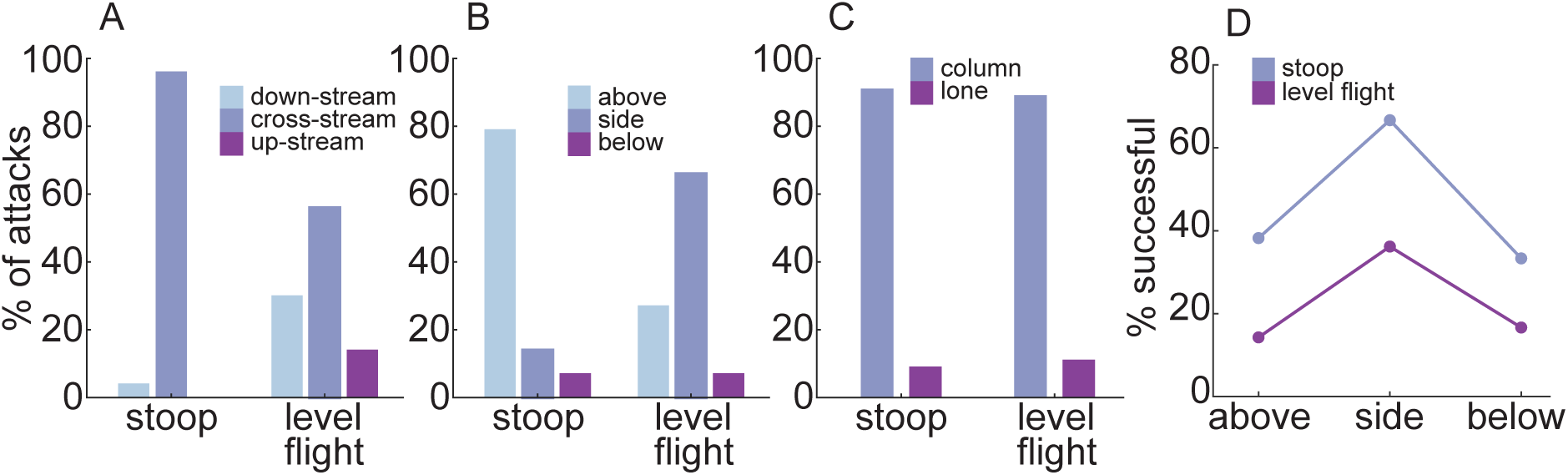
Behavioral interactions in bat-hunting Swainson’s hawks. (A-C) Proportions of attacks involving different categories of behavior in stooping versus level flight for (A)approach direction; (B) grab direction; (C) targeting strategy. (D) Interaction plot showing that using either a stooping dive or a rolling grab maneuver from the side increases catch successindependently.

### F. Hawks favor alternative attack strategies that increase catch success

The vast majority of the *n* = 218 attacks that we were able to categorise by approach direction involved either a cross-stream (65.1% of attacks; CI: 58.6%, 71.2%; Fig. 3A) or downstream approach (24.3% of attacks; CI: 19.1%, 30.4%; Fig. 3A). Upstream approaches were infrequent (10.6% of attacks; CI: 7.1%, 15.3%; Fig. 3A), which is consistent with the prediction that hawks should avoid attacking bats frontally to avoid their visual and acoustic gaze, although we found no statistical evidence that approach direction influenced the odds of capture. Approximately one-fifth of the *n* = 202 attacks that we were able to categorise fully involved stooping (21.3% of attacks; CI: 16.2%, 27.4%; Fig. 3A). Almost all of these *n* = 43 stoops involved a cross-stream approach (95.3% of stoops; CI: 84.5%, 98.7%; Fig. 4A), and most involved a pitching grab maneuver from above (79.1% of stoops; CI: 64.8%, 88.6%; Fig. 4B). In contrast, the majority of the *n* = 159 attacks made in level flight involved a rolling grab maneuver from the side (66.0% of attacks; CI: 58.4%, 72.9%; Fig. 4B). Hence, the great majority of the *n* = 202 attacks that we were able to categorise fully involved either a stoop or a rolling grab maneuver (73.3% of attacks; CI: 66.8%, 78.9%), these being the only two behavioral tactics that were associated with significantly improved capture odds. These two tactics were rarely used in combination, however (3.0% of attacks; CI: 1.4%, 6.3%). We found no evidence that the hawks modulated their attack strategy in relation to whether they were attacking a lone bat or the column, as target type was not significantly related to approach type (Fig. 4C), approach direction, or grab direction in the corresponding AR(1) logistic regression model (likelihood ratio tests at *α* = 0.05).

## 4. Discussion

### A. An adaptive account of the bats’ swarming behavior

The bats’ emergence in broad daylight is presumably driven by either: (i) a need to fly long distances to reach their feeding grounds, coupled with a need to time their arrival to coincide with the activity of their insect prey (Fenton et al. 1994), or (ii) a need to find water early, given the high daytime temperatures that can be experienced in their caves (Herreid 1963). Both challenges must be particularly acute at the Jornada Caves, which are located on a parched lava field 16km from the closest point on the Rio Grande, separated by desert. As a result of these constraints, the bats are exposed to intense predation by diurnal raptors on emergence, and it is not surprising that they return to the caves under cover of darkness, having met their daily needs for food and water. The 74 bats whose capture we have documented here through 64 focal follows made over 15 days constitute only ∼0.01% of the colony, but they represent a small fraction of the total catch over the observation period. Considering that the local population of hawks sustains its hunting activity over 4 to 5 months, it is probable that in order of magnitude terms ∼0.1% of the bat population falls victim to the hawks each year. Against such strong selection, it makes sense to ask whether the collective behavior that the bats exhibit on emergence reduces their individual risk of predation.

It was rare to see individual bats flying far from the column, and given that only ∼0.2% of the total population were classified as lone bats in sample images of their emergence, compared to ∼5% of those we recorded being caught (Section 2B), it is clear that the predation risk of lone bats must have been an order of magnitude higher than for bats in the column (Section 3B). In fact, some of the successful attacks that we categorised as being made on the column will actually have been attacks on lone bats silhouetted against it, leading us to underestimate systematically the odds that a captured bat was flying alone (*ô*_L|S_ = 0.048; CI: 0.016, 0.143). Furthermore, bats whose silhouettes overlapped within the column would have been counted as one, leading us to understate the swarm dilution factor as *D* > 500. It follows that our estimate of the relative risk of predation for lone bats, 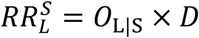, is expected to understate the adaptive benefits of column formation, such that 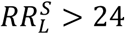. Bats maintaining column formation therefore benefit from a substantial reduction in their individual predation risk, so there should be strong selection in favor of mechanisms promoting aggregation on emergence. For example, swarming Brazilian free-tailed bats produce echolocation calls with different acoustic characteristics to foraging calls as they depart their caves in dense column formation, which has been hypothesized either to facilitate identification of spatial position within the group, or to serve in social communication (Gillam et al. 2010). Away from this central place, the expected encounter rate of lone predator and lone prey may be so low as to make the benefits of maintaining column formation negligible, or perhaps even negative given the high visibility and long detection range of the column. This may partly explain why the swarm retains a tight column formation on emergence, but loses its coherence away from the cave.

Capture rate per attack was not significantly different for bats flying alone or in the column, so we found no evidence that swarming bats benefitted from any net vigilance or confusion effect. However, as each attack on the column could involve up to three attempted grabs if the preceding grab(s) had been unsuccessful, the hawks had multiple opportunities to catch a bat when attacking the column. This could in principle have masked any effects of vigilance or confusion depressing the capture rate per grab for attacks on the column, but a binomial model of catch success for attacks involving multiple grabs provided no evidence that this was so (Section 3D). On the contrary, the capture rates that we recorded per attack are consistent with the null hypothesis that the hawks achieve the same success rate per grab in attacks involving multiple grabs at the column as in attacks involving a single grab at a lone bat. The adaptive benefits of swarming must therefore derive not from vigilance or confusion, but from dilution.

This is confirmed by relating the proportion of all observed attacks that were classified as attacks on lone bats (∼10%) to the overall proportion of bats classified as flying alone (∼0.2%), which shows that the attack risk of lone bats must have been an order of magnitude higher than that of bats in the column (Section 3C). Similar to our estimate of the relative predation risk, our estimate of the relative risk of attack for a lone bat, 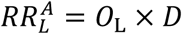 is also expected to understate the benefits of swarming, such that 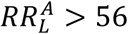 (CI: 37, 86). Dilution is therefore both necessary and sufficient to explain the higher survival rates of bats flying in the column.

### B. An adaptive account of the hawks’ hunting behavior

The attack abatement enjoyed by bats flying in the column is consistent with the expectation from classical selfish herd theory that lone individuals will have a larger domain of danger than those in a group, on the assumption that predators target whichever prey is closest to them as they attack (Duffield and Ioannou 2017; Hamilton 1971; Quinn and Cresswell 2006). Formal three-dimensional modelling of the chase dynamics (Brighton and Taylor 2019; Brighton et al. 2017) will be needed in the future to test whether the hawks target specific individuals on approach, or whether they simply grab at whichever bat happens to be closest, having broadly targeted the swarm (see also (Hein et al. 2020)). Certainly, our observations provide no evidence that the hawks adopt systematically different attack strategies against lone bats versus bats flying in the column.

Our observations provide the first empirical evidence that stooping attacks by an aerial predator enhances their catch success against agile prey, more than trebling the odds of success compared to initiating an attack in level flight (Section 3E; Fig. 4D). This result is qualitatively consistent with a recent physics-based simulation study, which found that the catch success of model falcons attacking agile model prey was maximised by initiating the attack from a high-speed, high-altitude dive (Mills et al. 2018). In these simulations, stooping enhanced catch success through: (i) the higher aerodynamic forces available for maneuvering at higher airspeed; (ii) the lower roll inertia present with the wings tucked; and (iii) the outright speed advantage conferred in a tail chase (Mills et al. 2018). The Swainson’s hawks that we studied almost always used a pitching grab maneuver to catch bats when stooping, and tended to use a cross-stream approach, rather than a tail chase. The mechanistic benefits of stooping in this case are therefore likely to be either: (i) the higher aerodynamic forces available for maneuvering at higher airspeed; or (ii) the element of surprise that stooping may confer.

Despite its tactical advantages, stooping was only used in 21% of the attacks that we observed (Fig. 3A). This might have been because having lost altitude in a stoop, it was then more efficient to make subsequent attacks in level flight, but 9 of the observed hunting bouts involved repeated stoops. More importantly, perhaps, attacks made from level flight could be just as successful as attacks made from stoops, provided that they involved a rolling grab maneuver (Fig. 4D). Other things being equal, this more than trebled the odds of catch success relative to the use of a pitching grab maneuver, consistent with the mechanistic advantages expected when banking into a turn (Mills et al. 2018). Overall, the great majority of the attacks that we observed involved a rolling grab maneuver (55% of attacks) and/or a stooping dive (21% of attacks), confirming that the hawks had a strong tendency to adopt their two most successful behavioral tactics (Section 3F). Moreover, as these tactics were rarely combined in a single attack (3% of attacks), and as they were deployed in quick succession on 17 of the observed hunting bouts, it would appear that they represent alternative and complementary attack strategies, rather than idiosyncratic behavioral traits of particular individuals.

### C. Behavioral interaction effects

Although the swarm was occasionally scattered by an attack, we did not see any clear evidence of the agitation waves that characterise coordinated evasive behavior in flocking birds (Procaccini et al. 2011). In the vast majority of cases, the only definite evasive behavior that we saw was a last-ditch attempt to avoid capture by an individual at immediate risk of being grabbed. In any case, given that 13% of attacks on lone bats ended in their capture, evading an attack is certainly much less effective than avoiding it altogether, given the likely magnitude of the attack abatement resulting from flying in the column (Section 4B). This may partly reflect the challenges of predator detection, which is a necessary condition for predator evasion. Bats sense their environment using echolocation and vision, which owing to their forward-facing eyes and sonar can result in blind zones above and behind the bat (Lima and O’Keefe 2013). The relative infrequency of upstream approaches that we observed in the hawks (Figs. 3A, 4A) may therefore be adaptive (Section 3F), since a downstream or cross-stream approach avoids placing the attacker within the primary visual and acoustic gaze of its target, whilst simultaneously reducing the demands on the attacker’s guidance and control (Mills et al. 2018). We found no evidence that the hawks modulated their hunting tactics according to whether they were attacking a bat flying alone or in the column.

### D. Avian predation of bats as a global selection pressure

Although there are numerous anecdotal reports of birds hunting bats (for review, see Table S1), there have been only a few systematic studies of this behavior to date (Black et al. 1979; Fenton et al. 1994; Lee and McCracken 2001; Roberts et al. 1997; Rodriguezduran and Lewis 1985). This is surprising given that bats comprise over 20% of all mammalian species (Lima and O’Keefe 2013), and occur globally in large colonies that are subject to intense depredation almost wherever they occur (Mikula et al. 2016). Owls are the most significant avian predators of bats in the temperate zones (Speakman 1991), but bat-hunting behavior has been documented in at least 237 species of diurnal birds worldwide, mainly from the families Accipitridae, Falconidae, and Corvidae (Mikula et al. 2016). Predation by diurnal birds is often cited as an evolutionary driver of nocturnality in bats (Mikula et al. 2016; Speakman 1991), but empirical evidence is limited, and surprisingly little is known of the underlying selection pressures (Lima and O’Keefe 2013). In fact, although attacks by raptors on emerging columns of bats were mooted as an example of marginal predation in Hamilton’s seminal paper on the selfish herd (Hamilton 1971), only one previous study has analysed the behavior formally from the perspective of group living (Fenton et al. 1994). This study of African raptors concluded that the straightforward numerical dilution of individual predation risk expected with increasing colony size is modified by: (i) the tendency of raptors to hunt at large colonies; (ii) the duration of emergence, which sets the time available for hunting in small colonies; and (iii) the time of emergence, which may limit the time available for hunting in very large colonies (Fenton et al. 1994).

Most avian predation of bats is opportunistic, with bats forming part of a broader diet (Mikula et al. 2016). Swainson’s hawks (*Buteo swainsoni*) are opportunistic diurnal hunters that consume a wide variety of mainly terrestrial mammals, reptiles, birds, and insects (Bednarz 1988), but have been recorded hunting swarming bats in three locations to date (Baker 1962; Cartron 2010; Harden 1972). Other raptors that hunt bats opportunistically do so using a broad range of behaviors, including stooping or swooping, level pursuit, and perch hunting (Table S1). We observed a similarly flexible range of behaviors in Swainson’s hawks, but we did not observe them perch hunting. The high overall catch success that we observed (31% of attacks resulting in capture) is not unusual in comparison to the 9 other species of diurnal raptor for which ≥15 capture outcomes have been recorded previously, with only red-tailed hawks (*Buteo jamaicensis*) and bat hawks (*Macheiramphus alcinus*) having significantly higher documented catch success (Fig. 5). Catch success varies greatly according to local conditions, and the exceptionally high catch success reported from red-tailed hawks opportunistically hunting another massive maternal colony of Brazilian free-tailed bats (68% of attacks resulting in capture) is thought to have been skewed by the emergence of juvenile bats midway through the observation period (Lee and Kuo 2001). Our own observations of Swainson’s hawks were made prior to the emergence of any juvenile bats, so will not have been skewed in this way.

**Figure 5.**
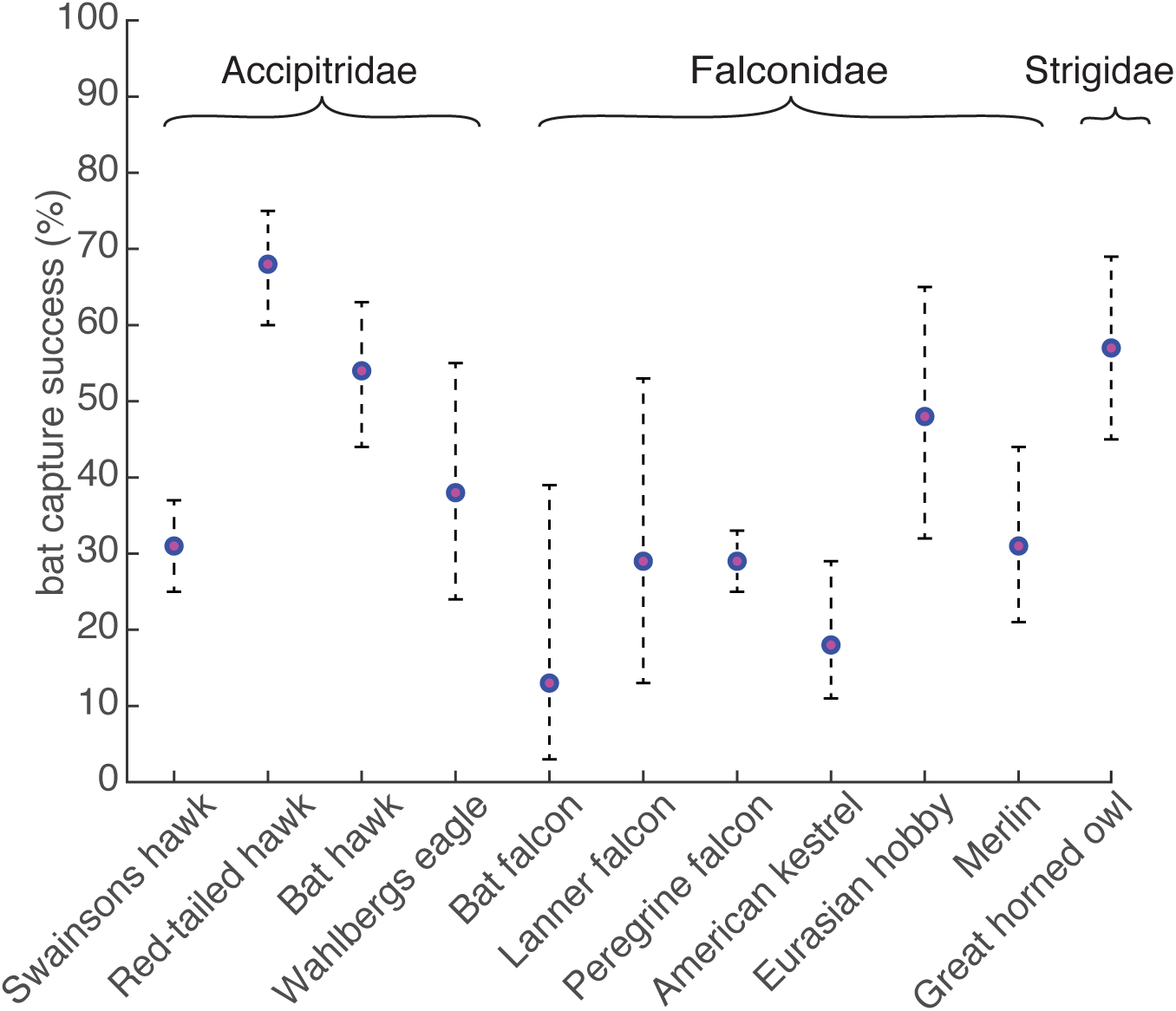
Recorded catch success against bats of 10 species of bird with ≥ 15 attacks recorded in the literature (Table S1), compared with the results of this study for Swainson’s hawks. Error bars display Agresti-Coull 95% CIs.

The very high catch success reported from bat hawks (54% of attacks resulting in capture) is perhaps more routine – this being the only species of bird that is considered morphologically, behaviorally, and ecologically specialized for hunting bats (Jones et al. 2012). Some of the opportunistic hunting behaviors that we observed in Swainson’s hawks parallel the specialist hunting behaviors of bat hawks, which are reported to intercept bats emerging from their roost in a series of back-and-forth flights (Black et al. 1979) (see also (Auburn 1987)), similar to the cross-stream approaches that were prevalent in Swainson’s hawks attacking the column (Fig. 3A). Likewise, the Swainson’s hawks that we observed consumed their prey on the wing (see also (Cartron 2010; Harden 1972)), paralleling a well-known behavioral adaptation of bat hawks (Ansell 1969; Auburn 1987; Ballance 1981; Black et al. 1979) to which their unusually large gape is supposed to be a morphological adaptation (Jones et al. 2012). Feeding in flight is an unusual behavior among raptors, except those like the Eurasian hobby (*Falco subbuteo*) that specialise on smaller aerial prey. Aerial feeding has also been recorded in a peregrine falcon (*Falco peregrinus*) hunting swarming bats (Sprunt 1951), so we suggest that this behavior is particularly adaptive during mass emergence, when the glut of prey makes it possible to capture multiple bats in a limited time window.

To conclude, colonial bats across the globe run a daily gauntlet against predatory birds as they emerge from or return to their roosts. The localisation, repeatability, and predictability of this behavior makes it an outstanding model system for studying predator-prey interactions and group dynamics in their natural ecological context. Moreover, as independent species assemblages of predators and prey perform essentially similar behaviors at different locations around the world, this system also lends itself to comparative study.

## Acknowledgements

We thank Turner Enterprises for access to and housing at the field location. We thank Lucy Larkman, Christian Harding, and Paul Domski for fieldwork assistance, and Yanqing Fu for the aerial photo of the cave.

## Authors’ contributions

CB, LK, and GT conceived the study. All authors contributed field observations. LZ and CB analysed video data. CB, LZ and GT performed statistical analysis. CB and GT wrote the paper, with input from LZ and LK. All authors commented on and approved the final version of the manuscript.

## Data and Code

Data and code implementing the statistical analysis are available through figshare: https://doi.org/10.6084/m9.figshare.11823393. The original video data are archived institutionally, and will be made available by the corresponding author upon reasonable request. Example videos are provided as Movie S1, and are available through figshare.

