## Supplementary Figures, Tables, and Legends for "Aerial attack strategies of bat-hunting hawks, and the dilution effect of swarming"

This file contains:

Figures S1-S2

Tables S1-S3

Supplementary References supporting Table S1

Legend for Movie S1

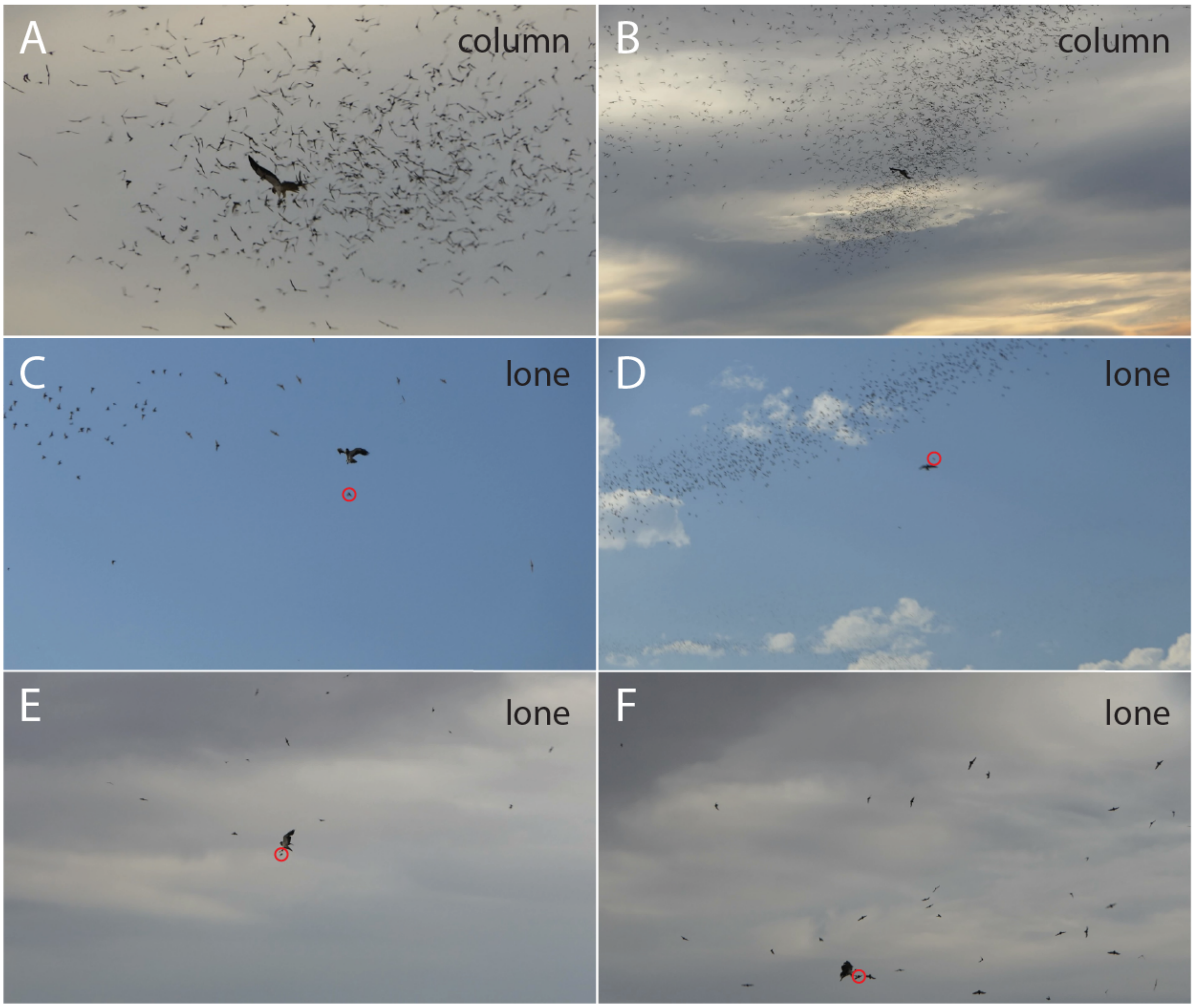


**Figure S1.** Video frames showing examples of attacks on lone bats and the column. (A,B) Attacks on the column of bats, defined as an attack on one or more bats within a cohesive group of individuals all flying in the same general direction. (C-E) Attacks on a lone bat (circled red), defined as an attack on an individual that appeared to be flying at least 1m from the edge of the column, and typically in a different direction to the swarm. (F) If an attack occurred in a volume containing many bats, but with no coherent flight direction, then this was also categorised as an attack on a lone bat, rather than as an attack on the swarm.

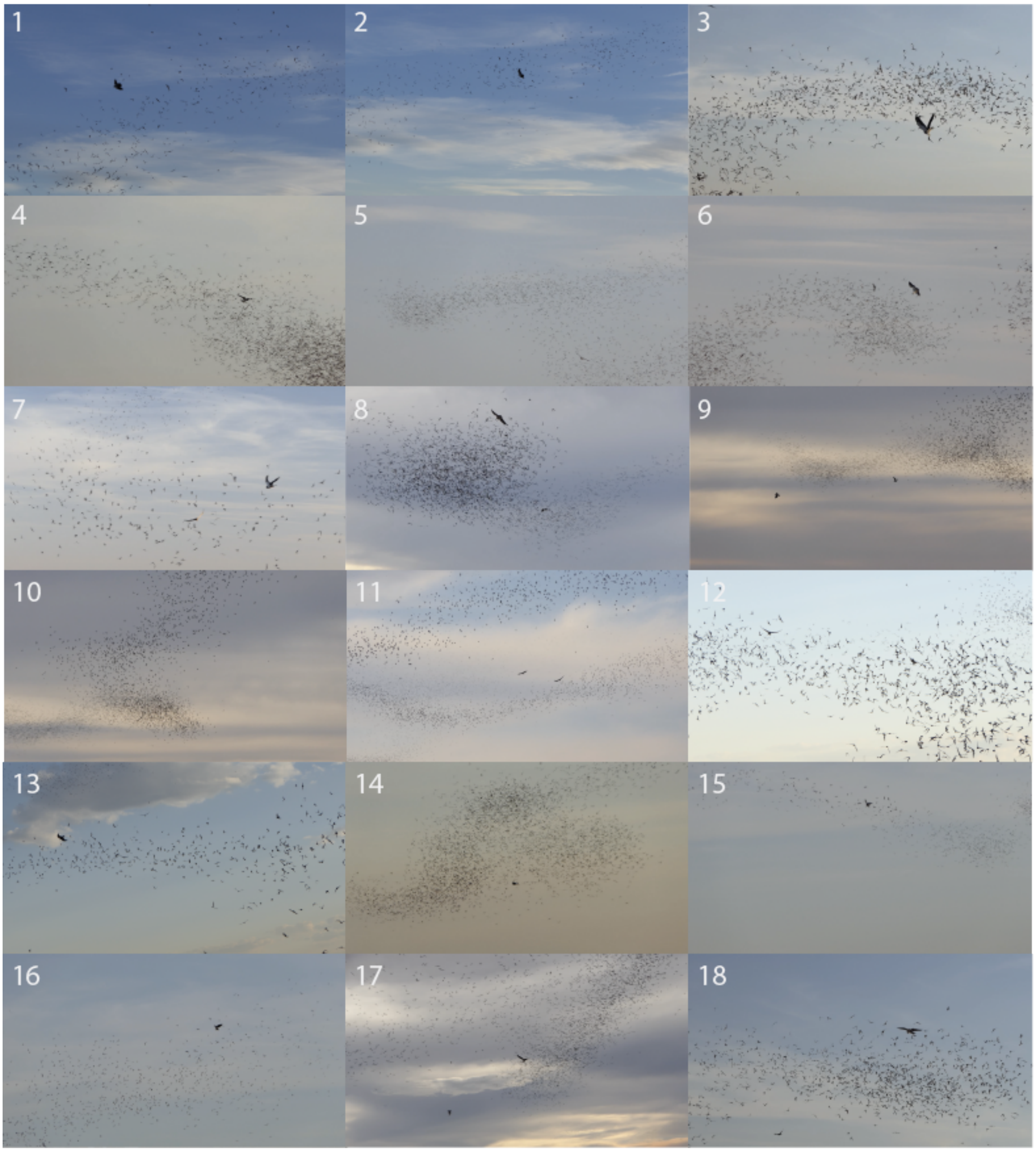


**Figure S2** Video frames used to estimate the proportion of bats meeting the criteria for classification as lone bats. These frames were chosen as meeting the following criteria: (i) each frame recorded during a separate attack; (ii) camera zoomed out and in focus; (iii) bats close enough to see their wings; (iv) background composed entirely of sky. See Table S3 for the results of this analysis.

| common name | scientific name | stoop or swoop | tail-chase | perch-hunting | hunting lone bat | hunting swarm | dawn | daytime | dusk | cooperative hunting | successes | observations | catch success | lower 95% CI | upper 95% CI | references |
|---|---|---|---|---|---|---|---|---|---|---|---|---|---|---|---|---|
| **Accipitriformes** | | | | | | | | | | | | | | | | |
| Bat hawk | *Macheiramphus alcinus* | y | y | | y | y | | | y | | 59 | 110 | 54% | 44% | 63% | [1-5] |
| Wahlberg's eagle | *Hieraaetus wahlbergi* | y | | | | y | | | y | | 13 | 34 | 38% | 24% | 55% | [6, 7] |
| Mississippi kite | *Ictinia mississippiensis* | y | | | | y | y | | y | | - | - | | | | [8] |
| Double-toothed kite | *Harpagus bidentatus* | | | y | y | y | | y | y | | 3 | 4 | | | | [9, 10] |
| Northern harrier | *Circus hudsonius* | y | | | | y | | | | | 0 | - | | | | [11] |
| Gabar goshawk | *Micronisus gabar* | y | | y | y | | | y | y | | 1 | 2 | | | | [12, 13] |
| African goshawk | *Accipiter tachiro* | | y | y | | y | | | y | | 11 | 14 | | | | [6, 14-16] |
| Cooper's hawk | *Accipiter cooperii* | | y | | y | y | | | y | | 1 | 1 | | | | [10, 17] |
| Eurasian sparrowhawk | *Accipiter nisus* | | y | | y | y | | y | y | | 0 | 10 | | | | [18, 19] |
| Shikra | *Accipiter badius* | y | y | y | | y | | | y | | - | - | | | | [20] |
| Harris's hawk | *Parabuteo unicinctus* | | | y | y | | | | y | | 2 | - | | | | [21] |
| Red-tailed hawk | *Buteo jamaicensis* | y | | y | | y | y | | y | | 96 | 142 | 68% | 60% | 75% | [10, 11, 22-24] |
| Swainson's hawk | *Buteo swainsoni* | y | y | | | y | | | y | | 7 | 8 | | | | [10, 23] |
| **Falconiformes** | | | | | | | | | | | | | | | | |
| Lanner falcon | *Falco biarmicus* | y | y | | y | y | | | y | | 5 | 17 | 29% | 13% | 53% | [25-27] |
| Merlin | *Falco columbarius* | y | y | y | | y | | | y | | 18 | 58 | 31% | 21% | 44% | [28, 29] |
| Dickinson's kestrel | *Falco dickinsoni* | y | | y | y | y | | | y | | 10 | - | | | | [30, 31] |
| Red-headed falcon | *Falco chicquera* | | y | y | | y | | | y | | ≥33 | - | | | | [32-34] |
| Peregrine falcon | *Falco peregrinus* | y | y | y | y | y | y | y | y | | 141 | 490 | 29% | 25% | 33% | [24, 35-38] |
| Bat falcon | *Falco rufigularis* | y | | y | y | | y | y | y | | 2 | 15 | 13% | 3% | 39% | [39, 40] |
| American kestrel | *Falco sparverius* | y | | y | y | y | y | y | y | | 12 | 66 | 18% | 11% | 29% | [10, 23, 41-43] |
| Eurasian hobby | *Falco subbuteo* | y | y | y | y | y | y | | y | | 15 | 31 | 48% | 32% | 65% | [6, 44-47] |
| Australian hobby | *Falco longipennis* | y | y | | | y | | | y | | 3 | 4 | | | | [48] |
| Sooty falcon | *Falco concolor* | | y | | y | | | | y | | 4 | 0 | | | | [49] |
| Common kestrel | *Falco tinnunculus* | y | | y | y | y | y | y | y | | 137 | - | | | | [50-52] |
| Lesser kestrel | *Falco naumanni* | y | y | | y | y | y | y | y | | 0 | ≥5 | | | | [67] |
| Australian kestrel | *Falco cenchroides* | y | | y | y | | | y | | | 1 | - | | | | [53] |
| Rock kestrel | *Falco rupicolus* | y | | | | y | | | y | | 1 | 0 | | | | [54] |
| **Strigiformes** | | | | | | | | | | | | | | | | |
| Barn owl | *Tyto alba* | y | | y | | y | | | y | | 3 | 4 | | | | [11, 42] |
| Great horned owl | *Bubo virginianus* | | | y | | y | | | y | | 36 | 63 | 57% | 45% | 69% | [55-57] |
| Northern long-eared owl | *Otus asio* | | | y | | y | | | y | | ≤5 | 12 | | | | [58] |
| **Passeriformes** | | | | | | | | | | | | | | | | |
| Carrion crow | *Corvus corone* | | | y | y | | y | | | | 1 | 2 | | | | [59] |
| Rook | *Corvus frugilegus* | | y | y | y | | | | y | | 0 | 3 | | | | [60] |
| American crow | *Corvus brachyrhynchos* | y | y | | y | | | y | y | y | 2 | 5 | | | | [61, 62] |
| Large-billed crow | *Corvus macrorhynchos* | y | | | | y | y | y | | y | >>1 | - | | | | [63] |
| Black-billed crow | *Pica hudsonia* | | y | | y | | | y | | | 1 | 1 | | | | [64] |
| Great grey shrike | *Lanius excurbitor* | | | y | | y | | | y | | 0 | ≥2 | | | | [65] |
| **Charadriiformes** | | | | | | | | | | | | | | | | |
| European herring gull | *Larus argentatus* | | y | | y | | | | | | 1 | 1 | | | | [66] |

**Table S1**. Summary of the results of previous studies recording observations of bat-hunting behaviours in birds. Each of the various categories of hunting behaviours is scored “y” if recorded at least once in that species, and left blank otherwise. A successful attack is defined as an attack in which a bat was caught, regardless of whether it was then eaten. Any study that only reported the number of successful attacks without also stating the total number of attacks observed was excluded from the calculation of catch success. Reported confidence intervals (CIs) are approximate 95% CIs calculated using the Agresti-Coull method after pooling all of the data.

| Date | attempted attacks | duration of focal follow (s) | approach type | | approach direction | | | targeting strategy | | grab direction | | | bat captured | | wind speed (mph) | wind direction (degs) |
|---|---|---|---|---|---|---|---|---|---|---|---|---|---|---|---|---|
| | | | stooping dive | level flight | down-stream | cross-stream | up-stream | column of bats | lone bat | above | side | below | yes | no | | |
| 01/06/2018 | 12 | 212 | 4 | 8 | 3 | 7 | 2 | 12 | 0 | 4 | 7 | 1 | 2 | 10 | 4.1 | 210 |
| 01/06/2018 | 7 | 208 | 0 | 7 | 3 | NaN | 2 | 6 | NaN | NaN | 4 | | 1 | 6 | 4.1 | 210 |
| 01/06/2018 | 4 | 69 | NaN | 3 | 1 | NaN | NaN | 3 | 1 | NaN | NaN | NaN | 0 | 4 | 4.1 | 210 |
| 01/06/2018 | 3 | 75 | 1 | 2 | 2 | 1 | NaN | 3 | NaN | 1 | NaN | NaN | 0 | 3 | 4.1 | 210 |
| 01/06/2018 | 8 | 180 | 1 | 7 | 3 | 5 | 0 | 8 | 0 | 1 | 6 | 1 | 2 | 6 | 4.1 | 210 |
| 02/06/2018 | 9 | 552 | 4 | 5 | NaN | 8 | NaN | 8 | 1 | 3 | 1 | NaN | 1 | 8 | 3.9 | 230 |
| 02/06/2018 | 8 | 150 | 0 | 8 | 3 | 2 | NaN | 8 | 0 | NaN | 5 | NaN | 1 | 7 | 3.9 | 230 |
| 02/06/2018 | 2 | 36 | 0 | 2 | 1 | NaN | NaN | 2 | 0 | NaN | 1 | NaN | 0 | 2 | 3.9 | 230 |
| 02/06/2018 | 2 | 114 | 2 | 0 | 0 | 2 | 0 | 1 | 1 | 1 | 1 | 0 | 1 | 1 | 3.9 | 230 |
| 02/06/2018 | 3 | 297 | 1 | 2 | 0 | 3 | 0 | 2 | NaN | NaN | 2 | NaN | 3 | 0 | 3.9 | 230 |
| 04/06/2018 | 3 | 175 | 1 | 2 | 1 | 2 | 0 | 3 | 0 | 2 | 1 | 0 | 3 | 0 | 4.1 | 230 |
| 04/06/2018 | 7 | 495 | 4 | 3 | 1 | 6 | 0 | 6 | 1 | 1 | NaN | 6 | 4 | 3 | 4.1 | 230 |
| 04/06/2018 | 2 | 97 | 0 | 2 | 0 | 1 | 1 | 1 | 1 | 1 | 1 | 0 | 1 | 1 | 4.1 | 230 |
| 04/06/2018 | 5 | 215 | 2 | 2 | NaN | 3 | 1 | 4 | NaN | 2 | 1 | 1 | 2 | 3 | 4.1 | 230 |
| 04/06/2018 | 4 | 286 | 1 | 1 | NaN | 1 | | 1 | NaN | NaN | NaN | NaN | 2 | 2 | 4.1 | 230 |
| 04/06/2018 | 8 | 28 | 0 | 8 | 0 | 0 | 8 | 8 | 0 | 5 | 3 | 0 | 0 | 8 | 4.1 | 230 |
| 04/06/2018 | 3 | 78 | 0 | 3 | 1 | 2 | 0 | 3 | 0 | 0 | 3 | 0 | 1 | 2 | 4.1 | 230 |
| 04/06/2018 | 2 | 32 | 0 | 2 | 1 | 1 | 0 | 2 | 0 | 0 | 2 | 0 | 1 | 1 | 4.1 | 230 |
| 04/06/2018 | 2 | 164 | 1 | 1 | 0 | 2 | 0 | 2 | 0 | 1 | 1 | 0 | 2 | 0 | 4.1 | 230 |
| 04/06/2018 | 5 | 171 | 0 | 5 | 2 | 3 | 0 | 5 | 0 | 2 | 3 | 0 | 1 | 4 | 4.1 | 230 |
| 05/06/2018 | 15 | 726 | 2 | 13 | 3 | 11 | 1 | 11 | 4 | 5 | 9 | 1 | 5 | 10 | 5.3 | 220 |
| 05/06/2018 | 2 | 28 | 2 | 0 | 0 | 2 | 0 | 2 | 0 | 2 | 0 | 0 | 1 | 1 | 5.3 | 220 |
| 05/06/2018 | 2 | 155 | 1 | 1 | 2 | 0 | 0 | 2 | 0 | 1 | NaN | NaN | 2 | 0 | 5.3 | 220 |
| 05/06/2018 | 1 | 47 | 0 | 1 | 0 | 1 | 0 | 1 | 0 | 0 | 1 | 0 | 1 | 0 | 5.3 | 220 |
| 05/06/2018 | 1 | 90 | 0 | 1 | 1 | 0 | 0 | 1 | 0 | 0 | 1 | 0 | 1 | 0 | 5.3 | 220 |
| 05/06/2018 | 4 | 259 | 1 | 2 | NaN | 3 | NaN | 3 | NaN | 1 | 2 | NaN | 3 | 1 | 5.3 | 220 |
| 05/06/2018 | 1 | 105 | 0 | 1 | 0 | 0 | 1 | 1 | 0 | 0 | 1 | 0 | 1 | 0 | 5.3 | 220 |
| 05/06/2018 | 1 | 205 | 0 | 1 | 0 | 1 | 0 | 1 | 0 | 0 | 1 | 0 | 1 | 0 | 5.3 | 220 |
| 06/06/2018 | 1 | 160 | NaN | NaN | NaN | NaN | NaN | NaN | NaN | NaN | NaN | NaN | 1 | 0 | 21 | 330 |
| 06/06/2018 | 1 | 75 | NaN | NaN | NaN | NaN | NaN | NaN | NaN | NaN | NaN | NaN | 1 | 0 | 21 | 330 |
| 07/06/2018 | 3 | 82 | 0 | 3 | 3 | 0 | 0 | 2 | 1 | 0 | 3 | 0 | 1 | 2 | 6.1 | 310 |
| 07/06/2018 | 3 | 120 | NaN | 2 | 1 | 1 | | 3 | 0 | | 1 | 1 | 1 | 2 | 6.1 | 310 |
| 07/06/2018 | 7 | 20 | 0 | 7 | 3 | 3 | 1 | 7 | 0 | 2 | 5 | 0 | 1 | 6 | 6.1 | 310 |
| 10/06/2018 | 7 | 394 | 1 | 6 | 0 | 5 | 2 | 7 | 0 | 0 | 7 | 0 | 4 | 3 | 7.1 | 320 |
| 11/06/2018 | 3 | 66 | 1 | 2 | 1 | 2 | 0 | 3 | 0 | 2 | 1 | 0 | 1 | 2 | 9.4 | 350 |
| 11/06/2018 | 2 | 115 | NaN | 1 | NaN | 1 | NaN | 1 | NaN | NaN | 1 | NaN | 2 | 0 | 9.4 | 350 |
| 11/06/2018 | 3 | 49 | 0 | 3 | 0 | 3 | 0 | 3 | 0 | 0 | 2 | 1 | 0 | 3 | 9.4 | 350 |
| 12/06/2018 | 3 | 85 | 1 | 2 | 0 | 2 | 1 | 3 | 0 | 1 | 2 | 0 | 1 | 2 | 8.2 | 210 |
| 18/06/2018 | 3 | 36 | 0 | 3 | 0 | 3 | 0 | 3 | 0 | 1 | 2 | 0 | 2 | 1 | 9.1 | 150 |
| 18/06/2018 | 5 | 197 | 0 | 5 | 0 | 5 | 0 | 5 | 0 | 3 | 2 | 0 | 1 | 4 | 9.1 | 150 |
| 18/06/2018 | 2 | 40 | 0 | 2 | 1 | 1 | 0 | 2 | 0 | 1 | 1 | 0 | 0 | 2 | 9.1 | 150 |
| 18/06/2018 | 4 | 235 | 1 | 3 | 1 | 2 | 1 | 4 | 0 | 2 | 2 | 0 | 2 | 2 | 9.1 | 150 |
| 19/06/2018 | 3 | 180 | 1 | 2 | 0 | 3 | 0 | 3 | 0 | 1 | 1 | 1 | 1 | 2 | 9.1 | 140 |
| 19/06/2018 | 1 | 30 | 0 | 1 | 0 | 1 | 0 | 0 | 1 | 0 | 1 | 0 | 0 | 1 | 9.1 | 140 |
| 19/06/2018 | 1 | 107 | 0 | 1 | 0 | 1 | 0 | 1 | 0 | 0 | 1 | 0 | 0 | 1 | 9.1 | 140 |
| 19/06/2018 | 9 | 439 | 1 | 8 | 5 | 4 | 0 | 4 | 4 | 5 | 3 | NaN | 1 | 8 | 9.1 | 140 |
| 21/06/2018 | 2 | 201 | 1 | 1 | 0 | 2 | 0 | 2 | 0 | 1 | 1 | 0 | 2 | 0 | 2.4 | 120 |
| 21/06/2018 | 5 | 231 | 0 | 5 | 1 | 4 | 0 | 5 | 0 | 3 | 2 | 0 | 0 | 5 | 2.4 | 120 |
| 22/06/2018 | 1 | 30 | 0 | 1 | 0 | 1 | 0 | 1 | 0 | 0 | 1 | 0 | 0 | 1 | 8.9 | 220 |
| 22/06/2018 | 4 | 108 | 0 | 4 | 0 | 4 | 0 | 1 | 3 | 2 | 2 | 0 | 1 | 3 | 8.9 | 220 |
| 22/06/2018 | 6 | 75 | 0 | 6 | 1 | 5 | 0 | 6 | 0 | 2 | 3 | 1 | 0 | 6 | 8.9 | 220 |
| 22/06/2018 | 3 | 60 | 0 | 3 | NaN | 1 | NaN | 1 | 2 | 1 | 2 | 0 | 0 | 3 | 8.9 | 220 |
| 23/06/2018 | 1 | 119 | 1 | 0 | 0 | 1 | 0 | 1 | 0 | 1 | 0 | 0 | 1 | 0 | 10.1 | 140 |
| 23/06/2018 | 9 | 122 | 7 | 2 | 2 | 7 | 0 | 7 | 2 | 5 | 1 | NaN | 1 | 8 | 10.1 | 140 |
| 23/06/2018 | 2 | 240 | 1 | 1 | 1 | 1 | 0 | 2 | 0 | 1 | 1 | 0 | 1 | 1 | 10.1 | 140 |
| 23/06/2018 | 1 | 250 | 0 | 1 | 0 | 1 | 0 | 1 | 0 | 1 | 0 | 0 | 1 | 0 | 10.1 | 140 |
| 23/06/2018 | 3 | 120 | 1 | 2 | 0 | 3 | 0 | 3 | 0 | 1 | 2 | 0 | 0 | 3 | 10.1 | 140 |
| 23/06/2018 | 1 | 480 | 0 | 1 | 0 | 1 | 0 | 1 | 0 | 1 | 0 | 0 | 1 | 0 | 10.1 | 140 |
| 23/06/2018 | 7 | 108 | 3 | 4 | 1 | 4 | 2 | 6 | 1 | 6 | 1 | 0 | 1 | 6 | 10.1 | 140 |
| 23/06/2018 | 1 | 145 | 1 | 0 | 0 | 1 | 0 | 1 | 0 | 0 | 1 | 0 | 1 | 0 | 10.1 | 140 |
| 23/06/2018 | 1 | 123 | 0 | 1 | 1 | 0 | 0 | 1 | 0 | 0 | 1 | 0 | 1 | 0 | 10.1 | 140 |
| 24/06/2018 | 1 | 20 | 0 | 1 | 1 | 0 | 0 | 1 | 0 | 0 | 1 | 0 | 0 | 1 | 7.1 | 220 |
| 24/06/2018 | 2 | 60 | 0 | 2 | 0 | 2 | 0 | 2 | 0 | 1 | 0 | 1 | 0 | 2 | 11.6 | 320 |
| 24/06/2018 | 2 | 57 | 0 | 2 | 2 | 0 | 0 | 2 | 0 | 0 | 2 | 0 | 0 | 2 | 11.6 | 320 |
| **total** | **239** | **10228** | **49** | **181** | **53** | **142** | **23** | **205** | **23** | **77** | **113** | **15** | **74** | **165** | | |
| **% of total** | | | **21.3** | **78.7** | **24.3** | **65.1** | **10.6** | **89.9** | **10.1** | **37.6** | **55.1** | **7.3** | **31.0** | **69.0** | | |
| **95% CI (%)** | | | **16.2<br>27.4** | **72.6<br>83.8** | **19.1<br>30.4** | **58.6<br>71.1** | **7.1<br>15.3** | **85.3<br>93.2** | **6.8<br>14.7** | **31.2<br>44.4** | **48.3<br>61.8** | **4.5<br>11.7** | **25.4<br>37.1** | **62.9<br>74.6** | | |

**Table S2.** Classification of hawk attack flights against bats, from observational data. Orange highlights all observations made using the binocular follows, and grey shows observations made using video cameras. A 'NaN' denotes missing data. NB 'hawk no.' only highlights sequential hawks observed on a given night, but we could not identify these same hawks from day-to-day.

| frame | binarization threshold | total bat count | lone bat count | proportion of lone bats |
|---|---|---|---|---|
| 1 | 93 | 1137 | 7 | 0.62% |
| 2 | 110 | 868 | 3 | 0.35% |
| 3 | 133 | 1270 | 3 | 0.24% |
| 4 | 145 | 1558 | 2 | 0.13% |
| 5 | 140 | 2682 | 4 | 0.15% |
| 6 | 140 | 1874 | 3 | 0.16% |
| 7 | 154 | 702 | 2 | 0.28% |
| 8 | 131 | 2568 | 3 | 0.12% |
| 9 | 115 | 2168 | 2 | 0.09% |
| 10 | 125 | 2687 | 5 | 0.19% |
| 11 | 160 | 3576 | 4 | 0.11% |
| 12 | 213 | 1417 | 3 | 0.21% |
| 13 | 103 | 1234 | 7 | 0.57% |
| 14 | 132 | 3778 | 2 | 0.05% |
| 15 | 139 | 777 | 3 | 0.39% |
| 16 | 131 | 924 | 4 | 0.43% |
| 17 | 111 | 3404 | 4 | 0.12% |
| 18 | 119 | 1312 | 6 | 0.46% |
| **total** | | **33936** | **67** | **0.20%** |

**Table S3.** Proportion of bats meeting the criteria for classification as lone bats in the 18 video frames in Fig. S2. Bats meeting the criteria for classification as lone bats were counted individually in each frame, and compared to the total number of bats estimated using the object count function in Adobe Photoshop CC2019, after binarizing each image using a binarization threshold just sufficient to make the background sky entirely white. Because individuals with overlapping silhouettes are counted as a single object, this method results in a highly conservative estimate of the proportion of individuals meeting the criteria for classification as lone bats in each frame.

**Movie S1.** Example video sequences illustrating the behavioural classifications used to describe attacks by Swainson’s hawks on Brazilian free-tailed bats. See Box 1 for definitions of the various categories of behaviour. This video is compiled from clips recorded in a mixture of HD (1920×1080 pixels; 50 fps) and 4K UHD (3840×2160 pixels; 25 fps), uniformly downsampled to 1920×1080 pixels at 25 fps using H.264 compression.
