## Supplementary material for "Aerial attack strategies of bat-hunting hawks, and the dilution effect of swarming": Code S1, Data S1 and S2: CodeS1.R

```
#########################################################################################################
#This R-code implements the statistical analysis described in the paper:

#Aerial attack strategies of bat-hunting hawks, and the dilution effect of swarming
#Caroline H. Brighton, Lillias Zusi, Kathryn McGowan, Morgan Kinniry, Laura N. Kloepper, Graham K. Taylor
# 
#Department of Zoology, University of Oxford, South Parks Road, Oxford, OX1 3PS, UK.
#Department of Biology, Saint Mary’s College, Notre Dame, IN 46556, USA.

#The data needed to run this code are stored in the files DataS1.csv and DataS2.csv 
#available from figshare: https://doi.org/10.6084/m9.figshare.11823393
#########################################################################################################
# Preliminaries:
#########################################################################################################
setwd('~/Documents/Publications/in prep/Brighton et al bats/BehavEcol/')

# Load packages
library(PropCIs)
library(boot)
library(plyr)
#########################################################################################################
# Load data from DataS1.csv 
batData = read.table("DataS1.csv",header=TRUE, sep=",",)

# Declare variables and levels
batData$bout=factor(batData$bout)
batData$approachType=factor(batData$approachType,labels=c("stoop","level"))
batData$approachDirection=factor(batData$approachDirection,labels=c("down","cross","up"))
batData$targetType=factor(batData$targetType,labels=c("column","lone"))
batData$grabDirection=factor(batData$grabDirection,labels=c("above","side","below"))
batData$outcome=factor(batData$outcome,labels=c("success","failure"))
batData$moulting=factor(batData$moulting,labels=c("notmoulting","moulting"))

#########################################################################################################
# Load data from DataS2.csv 
boutData = read.table("DataS2.csv",header=TRUE, sep=",",)

#########################################################################################################
# Create a copy of the data with rows with missing values eliminated
batDataCut<-na.omit(batData)
# Set convenient reference levels for variables
batDataCut$outcome<-relevel(batDataCut$outcome, "failure")
batDataCut$approachType<-relevel(batDataCut$approachType, "level")
batDataCut$approachDirection<-relevel(batDataCut$approachDirection, "cross")
batDataCut$grabDirection<-relevel(batDataCut$grabDirection, "above")

#########################################################################################################
# Define total duration of observations (s)    
duration<-sum(boutData$duration)

#########################################################################################################
cat("Statistical analysis supporting the paper:","\n")
cat("\n")
cat("Aerial attack strategies of bat-hunting hawks, and the dilution effect of swarming","\n")
cat("By Caroline H. Brighton, Lillias Zusi, Kathryn McGowan, Morgan Kinniry, Laura N. Kloepper, Graham K. Taylor","\n")

#########################################################################################################
#Methods. 2B. Behavioral classification
#########################################################################################################
cat("\n")
cat("---------------------------------------------------------------------------------------------", "\n")  
cat("Methods Section: 2B. Behavioral classification","\n")
cat("---------------------------------------------------------------------------------------------", "\n")
cat("Total number of attacks:", length(batData$outcome),"\n")
cat("Of which, total number classified fully:", length(batDataCut$outcome),"\n")
cat("Total number of hunting bouts:", length(levels(batData$bout)), "\n")
cat("---------------------------------------------------------------------------------------------", "\n")
cat("\n")
#########################################################################################################
cat("\n")

#########################################################################################################
#Methods. 2C. Statistical analysis
#########################################################################################################
cat("---------------------------------------------------------------------------------------------", "\n")  
cat("Methods Section: 2C. Statistical analysis","\n")
cat("---------------------------------------------------------------------------------------------", "\n")
#########################################################################################################
# Create lagged variable for attack outcome for all attacks
batData$outcomeAR1[1]<-batData$outcome[length(batData$outcome)]
batData$outcomeAR1[2:length(batData$outcome)]<-batData$outcome[1:length(batData$outcome)-1]
batData$outcomeAR1=factor(batData$outcomeAR1,labels=c("success","failure"))

#########################################################################################################
# Identify only those attacks representing consecutive attacks from the same hunting bout
batData$boutAR1[1]<-batData$bout[length(batData$bout)]
batData$boutAR1[2:length(batData$bout)]<-batData$bout[1:length(batData$bout)-1]
sameBoutLagged<-batData$bout==batData$boutAR1

#########################################################################################################
# Calculate correlation between outcome of each attack and the outcome of the one immediately preceding it
cat("For the", length(batData$outcome[sameBoutLagged]), "pairs of consecutive attacks within hunting bouts:","\n")
ccfOutcomes<-ccf(batData$outcome[sameBoutLagged],batData$outcomeAR1[sameBoutLagged], lag.max=0)
r1<-ccfOutcomes$acf[1]
cat("1st-order autocorrelation coefficient for attack outcomes:", r1, "\n")
cat("95% CI for white noise autocorrelation function:", qnorm((1 - 0.95)/2)/sqrt(length(batData$outcome[sameBoutLagged])), qnorm((1 + 0.95)/2)/sqrt(length(batData$outcome[sameBoutLagged])),"\n")

#########################################################################################################
# Create lagged variable for attack outcome for all fully classified attacks
batDataCut$outcomeAR1[1]<-batDataCut$outcome[length(batDataCut$outcome)]
batDataCut$outcomeAR1[2:length(batDataCut$outcome)]<-batDataCut$outcome[1:length(batDataCut$outcome)-1]
batDataCut$outcomeAR1=factor(batDataCut$outcomeAR1,labels=c("success","failure"))
cat("\n")

#########################################################################################################
cat("Provisional model-order selection for pure autoregressive logistic regression model of attack outcome:")
print(ar.mle(batDataCut$outcome, order.max=5, AIC=TRUE))
cat("---------------------------------------------------------------------------------------------", "\n")
cat("\n")

#########################################################################################################
cat("\n")
#########################################################################################################
#Results 3A. Hawks hunting swarming bats achieve high catch success at high intensity
#########################################################################################################
cat("---------------------------------------------------------------------------------------------", "\n")
cat("Results Section 3A: Hawks hunting swarming bats achieve high catch success at high intensity","\n")
cat("---------------------------------------------------------------------------------------------", "\n")
cat("Number of attacks per bout:", "\n")
print(summary(boutData$attacks))
cat("\n")

#########################################################################################################
cat("Quasi-Poisson regression analysis of attack rate:","\n")
modelAttackRate<-glm(boutData$attacks~offset(log(boutData$duration)), family="quasipoisson"(link="log"))
cat("Mean attack rate (s^-1):", exp(modelAttackRate$coefficients), "\n")
CIAttackRate<-suppressMessages(confint(modelAttackRate))
cat("95% CI:", exp(CIAttackRate), "\n")
summaryModelAttackRate<-summary(modelAttackRate)
cat("Dispersion parameter:", summaryModelAttackRate$dispersion, "\n")
cat("\n") 

#########################################################################################################
table4<-table(batData$bout,batData$outcome)
P7<-sum(table4[,1]!=0)/length(table4[,1])
CI7<-scoreci(sum(table4[,1]!=0),length(table4[,1]),0.95)
cat("Proportion of bouts resulting in capture of one or more bats:","\n")
cat(P7, "; 95% CI:", CI7$conf.int[1], CI7$conf.int[2], "\n")
cat("\n")

#########################################################################################################
cat("Quasi-Poisson regression analysis of number of bats caught per bout:","\n")
tableSuccesses<-table(batData$bout,batData$outcome)
modelBatsCaughtPerBout<-glm(tableSuccesses[,1]~1, family="quasipoisson"(link="log"))
cat("Number of bats caught per bout:", exp(modelBatsCaughtPerBout$coefficients), "\n")
CIBatsCaughtPerBout<-suppressMessages(confint(modelBatsCaughtPerBout))
cat("95% CI:", exp(CIBatsCaughtPerBout), "\n")
summaryModelBatsCaughtPerBout<-summary(modelBatsCaughtPerBout)
cat("Dispersion parameter:", summaryModelBatsCaughtPerBout$dispersion, "\n")
cat("\n") 

#########################################################################################################
cat("Quasi-Poisson regression analysis of number of bats caught per attack:","\n")
modelBatsCaughtPerAtttack<-glm(tableSuccesses[,1]~offset(log(boutData$attacks)), family="quasipoisson"(link="log"))
cat("Number of bats caught per attack:", exp(modelBatsCaughtPerAtttack$coefficients), "\n")
CIBatsCaughtPerAtttack<-suppressMessages(confint(modelBatsCaughtPerAtttack))
cat("95% CI:", exp(CIBatsCaughtPerAtttack), "\n")
summaryModelBatsCaughtPerAtttack<-summary(modelBatsCaughtPerAtttack)
cat("Dispersion parameter:", summaryModelBatsCaughtPerAtttack$dispersion, "\n")

#########################################################################################################
# Bootstrap analysis for mean hunting efficiency by bout
# Defines function for calculating mean hunting efficiency by bout
byBout <- function(X,ind) {
  tableEfficiency<-table(X$bout[ind],X$outcome[ind])
  meanEfficiency<-mean(tableEfficiency[,1]/rowSums(tableEfficiency))}
# Bootstrapping
nresamples<-1000 #Increase to 1000,000 to replicate analysis in paper (slow)
stratifiedBootstrap <- boot(batData, statistic=byBout, R = nresamples, strata = batData$bout)
bootstrappedCI<-boot.ci(stratifiedBootstrap, conf=0.95, type="bca")
cat("\n") 
cat("Mean success rate by bout:", bootstrappedCI$t0, "; 95% CI*:", bootstrappedCI$bca[1,4:5], "\n")
cat("*",nresamples,"stratified resamples used to compute bootstrap confidence interval by bout","\n")
cat("---------------------------------------------------------------------------------------------", "\n")
cat("\n")

#########################################################################################################
cat("\n")
#########################################################################################################
#Results 3B. Flying in the column reduces predation risk in swarming bats
#########################################################################################################
cat("---------------------------------------------------------------------------------------------", "\n")
cat("Results Section 3B: Flying in the column reduces predation risk in swarming bats","\n")
cat("---------------------------------------------------------------------------------------------", "\n")
table3<-table(batData$targetType,batData$outcome)
cat("Number of bats caught:", sum(table3[,1]),"\n")
print(table3)
P6<-table3[2,1]/sum(table3[,1])
CI6<-scoreci(table3[2,1],sum(table3[,1]),0.95)
O1<-table3[2,1]/table3[1,1]
cat("\n")
cat("p(L|S):", P6,"; 95% CI:",CI6$conf.int[1],CI6$conf.int[2],"\n")
cat("Odds that a caught individual was a lone bat:", O1, "; 95% CI:",CI6$conf.int[1]/(1-CI6$conf.int[1]),CI6$conf.int[2]/(1-CI6$conf.int[2]),"\n")
cat("---------------------------------------------------------------------------------------------", "\n")
cat("\n")
#########################################################################################################
cat("\n")

#########################################################################################################
#Results 3C. Dilution, not vigilance or confusion, reduces predation risk in the column 
#########################################################################################################
cat("---------------------------------------------------------------------------------------------", "\n")
cat("Results Section 3C. Dilution, not vigilance or confusion, reduces predation risk in the column","\n")
cat("---------------------------------------------------------------------------------------------", "\n")
table2<-table(batData$targetType)
cat("Number of attacks categorised by target type:", sum(table2), "\n")
print(table3)
cat("\n")
cat("AR(1) logistic regression model for attack outcome on target type","\n")
modeloutcomeAR1<-glm(outcome~outcomeAR1+targetType,family=binomial,data=batData)
print(formula(modeloutcomeAR1))
cat("\n")
print(anova(modeloutcomeAR1, test = "LRT"))
cat("\n")

#########################################################################################################
cat("\n")
cat("Proportion of attacks on each target type:")
print(table2)
cat("\n")
P2<-table2[1]/sum(table2)
CI2<-scoreci(table2[1],sum(table2),0.95)
cat("p(C):", P2,"; 95% CI:",CI2$conf.int[1],CI2$conf.int[2],"\n")
P3<-table2[2]/sum(table2)
CI3<-scoreci(table2[2],sum(table2),0.95)
cat("p(L):", P3,"; 95% CI:",CI3$conf.int[1],CI3$conf.int[2],"\n")
O0<-P3/P2
cat("Odds that an attack was on a lone bat:", O0, "; 95% CI:",CI3$conf.int[1]/(1-CI3$conf.int[1]),CI3$conf.int[2]/(1-CI3$conf.int[2]),"\n")
cat("---------------------------------------------------------------------------------------------", "\n")
cat("\n")
#########################################################################################################
cat("\n")

#########################################################################################################
#Results 3D. Hawks attacking the column have multiple opportunities to grab a bat
#########################################################################################################
cat("---------------------------------------------------------------------------------------------", "\n")
cat("Results Section 3D. Hawks attacking the column have multiple opportunities to grab a bat","\n")
cat("---------------------------------------------------------------------------------------------", "\n")
cat("Number of attacks categorised by target type:", sum(table2), "\n")
print(table3<-table(batData$targetType,batData$outcome))
cat("\n")
P5<-table3[2,1]/sum(table3[2,])
CI5<-scoreci(table3[2,1],sum(table3[2,]),0.95)
cat("p(S|L):", P5,"; 95% CI:",CI5$conf.int[1],CI5$conf.int[2],"\n")
cat("Expected catch success in an attack involving up to 3 independent grabs at this success rate:", 1-(1-P5)^3,"\n")
cat("\n")
P4<-table3[1,1]/sum(table3[1,])
CI4<-scoreci(table3[1,1],sum(table3[1,]),0.95)
cat("p(S|C):", P4, "; 95% CI:",CI4$conf.int[1],CI4$conf.int[2],"\n")
cat("---------------------------------------------------------------------------------------------", "\n")
cat("\n")
#########################################################################################################
cat("\n")

#########################################################################################################
#Results 3E. Stoops and rolling grab maneuvers are associated with higher catch success
#########################################################################################################
cat("---------------------------------------------------------------------------------------------", "\n")
cat("Results Section 3E. Stoops and rolling grab maneuvers are associated with higher catch success","\n")
cat("---------------------------------------------------------------------------------------------", "\n")
cat("Full AR(1) logistic regression model for attack outcome:","\n")
modeloutcomeAR1<-glm(outcome~outcomeAR1+targetType+approachType+grabDirection+approachDirection+windSpeed+timeToSunset,family=binomial,data=batDataCut)
print(summary(modeloutcomeAR1))
cat("\n")

#########################################################################################################
cat("Reduced AR(1) logistic regression model for attack outcome retaining only significant terms:","\n")
modeloutcomeAR1reduced<-glm(outcome~outcomeAR1+approachType+grabDirection,family=binomial,data=batDataCut)
print(summary(modeloutcomeAR1reduced))
cat("\n")

#########################################################################################################
cat("Full AR(1) logistic regression model for attack outcome:","\n")
print(modeloutcomeAR1$formula)
cat("AIC:", modeloutcomeAR1$aic, "\n")
cat("Delta AIC for full AR(1) model against reduced AR(1) model", modeloutcomeAR1$aic-modeloutcomeAR1reduced$aic, "\n")
cat("\n")

#########################################################################################################
modeloutcomeAR1null<-glm(outcome~outcomeAR1,family=binomial,data=batDataCut)
cat("Pure AR(1) logistic regression model for attack outcome:", "\n")
print(modeloutcomeAR1null$formula)
cat("AIC:", modeloutcomeAR1null$aic, "\n")
cat("Delta AIC for pure AR(1) model against reduced AR(1) model", modeloutcomeAR1null$aic-modeloutcomeAR1reduced$aic, "\n")
cat("\n")

#########################################################################################################
modeloutcomeAR1interaction<-glm(outcome~outcomeAR1+approachType+grabDirection+approachType*grabDirection,family=binomial,data=batDataCut)
cat("Reduced AR(1) logistic regression model wth interaction term:","\n")
print(modeloutcomeAR1interaction$formula)
cat("AIC:", modeloutcomeAR1interaction$aic, "\n")
cat("Delta AIC for AR(1) model with interaction term against reduced AR(1) model", modeloutcomeAR1interaction$aic-modeloutcomeAR1reduced$aic, "\n")
cat("\n")

#########################################################################################################
cat("\n")
cat("Exponentiated coefficients and 95% CIs from reduced AR(1) model:","\n")
CIAR1<-suppressMessages(confint(modeloutcomeAR1reduced, level = 0.95))
cat("\n")
print(exp(modeloutcomeAR1reduced$coefficients[3]))
print(exp(CIAR1[3,]))
cat("\n")
print(exp(modeloutcomeAR1reduced$coefficients[4]))
print(exp(CIAR1[4,]))
cat("\n")
cat("---------------------------------------------------------------------------------------------", "\n")
cat("\n")
#########################################################################################################
cat("\n")

#########################################################################################################
#Results 3F. Hawks favor alternative attack strategies that increase catch success
#########################################################################################################
cat("---------------------------------------------------------------------------------------------", "\n")
cat("Results Section 3F. Hawks favor alternative attack strategies that increase catch success","\n")
cat("---------------------------------------------------------------------------------------------", "\n")
table5<-table(batData$approachDirection)
cat("Number of attacks categorised by approach direction:", sum(table5), "\n") 
print(table5)
cat("\n")

#########################################################################################################
P11<-table5[2]/sum(table5)
CI11<-scoreci(table5[2],sum(table5),0.95)
cat("Proportion of cross-stream approaches", P11,"; 95% CI:",CI11$conf.int[1],CI11$conf.int[2],"\n")
P10<-table5[1]/sum(table5)
CI10<-scoreci(table5[1],sum(table5),0.95)
cat("Proportion of downstream approaches:", P10,"; 95% CI:",CI10$conf.int[1],CI10$conf.int[2],"\n")
P12<-table5[3]/sum(table5)
CI12<-scoreci(table5[3],sum(table5),0.95)
cat("Proportion of upstream approaches", P12,"; 95% CI:",CI12$conf.int[1],CI12$conf.int[2],"\n")
cat("\n")

#########################################################################################################
cat("\n")
cat("Number of attacks categorised fully:", length(batDataCut$outcome), "\n")
cat("\n")
print(combos<-count(batDataCut,c("approachType","approachDirection","grabDirection")))
P13<-combos[11,4]/length(batDataCut$outcome)
CI13<-scoreci(combos[11,4],length(batDataCut$outcome), 0.95)
P14<-(sum(combos[10:13,4])+combos[2,4]+combos[5,4]+combos[8,4])/length(batDataCut$outcome)
CI14<-scoreci((sum(combos[10:13,4])+combos[2,4]+combos[5,4]+combos[8,4]),length(batDataCut$outcome), 0.95)
P15<-(sum(combos[10:13,4]))/length(batDataCut$outcome)
CI15<-scoreci((sum(combos[10:13,4])),length(batDataCut$outcome), 0.95)
P18<-(sum(combos[10:12,4]))/(sum(combos[10:13,4]))
CI18<-scoreci(sum(combos[10:12,4]),sum(combos[10:13,4]), 0.95)
P19<-(combos[10,4]+combos[13,4])/(sum(combos[10:13,4]))
CI19<-scoreci((combos[10,4]+combos[13,4]),sum(combos[10:13,4]), 0.95)
P20<-(combos[2,4]+combos[5,4]+combos[8,4])/(sum(combos[1:9,4]))
CI20<-scoreci((combos[2,4]+combos[5,4]+combos[8,4]),sum(combos[1:9,4]), 0.95)
cat("\n")
cat("Proportion of fully-classified attacks involving stooping:",P15, "95% CI:",CI15$conf.int[1],CI15$conf.int[2],"\n")
cat("\n")
cat("Number of fully-classified attacks involving stooping::", sum(combos[10:13,4]), "\n")
cat("Proportion of stoops involving a cross-stream approach:",P18, "95% CI:",CI18$conf.int[1],CI18$conf.int[2],"\n")
cat("Proportion of stoops involving a pitching grab from above:", P19, "95% CI:",CI19$conf.int[1],CI19$conf.int[2],"\n")
cat("\n")
cat("Number of fully-classified attacks involving level flight:", sum(combos[1:9,4]), "\n")
cat("Proportion of level attacks involving a cross-stream approach:", P20, "95% CI:",CI20$conf.int[1],CI20$conf.int[2],"\n")
cat("\n")
cat("Proportion of fully-classified attacks involving stooping OR an attack from from the side:",P14, "95% CI:",CI14$conf.int[1],CI14$conf.int[2],"\n")
cat("Proportion of attacks involving stooping AND an attack from the side:",P13, "95% CI:",CI13$conf.int[1],CI13$conf.int[2],"\n")
cat("\n")
cat("\n")

#########################################################################################################
# Create lagged variable for target type for all fully classified attacks
batDataCut$targetTypeAR1[1]<-batDataCut$targetType[length(batDataCut$targetType)]
batDataCut$targetTypeAR1[2:length(batDataCut$targetType)]<-batDataCut$targetType[1:length(batDataCut$targetType)-1]
batDataCut$targetTypeAR1=factor(batDataCut$targetTypeAR1,labels=c("column","lone"))

#########################################################################################################
cat("Full AR(1) autoregressive logistic regression model for target type:", "\n")
cat("\n")
modeltargetTypeAR1<-glm(targetType~targetTypeAR1+approachType+approachDirection+grabDirection,family=binomial,data=batDataCut)
print(summary(modeltargetTypeAR1))
print(anova(modeltargetTypeAR1,test="LRT"))

#########################################################################################################
cat("---------------------------------------------------------------------------------------------", "\n")

#END OF SCRIPT
```
